# Expression of Tim-3 drives naïve Treg to an effector-like state with enhanced suppressive activity

**DOI:** 10.1101/2020.07.31.230714

**Authors:** Hridesh Banerjee, Hector Nieves-Rosado, Aditi Kulkarni, Benjamin Murter, Uma R. Chandran, Alexander Chang, Andrea L. Szymczak-Workman, Lazar Vujanovic, Robert L. Ferris, Lawrence P. Kane

## Abstract

Regulatory T cells (Treg) are critical mediators of self-tolerance but can also limit effective anti-tumor immunity. We and others previously reported that 40-60% percent of Treg-infiltrating head and neck cancer (HNC) and other tumors highly express Tim-3, compared with about 5% in lymphoid organs. Tumor-infiltrating Tim-3^+^ Treg also have enhanced suppressive function and display a more effector-like phenotype. Using a novel mouse model with cell type-specific Tim-3 expression, we show here that expression of Tim-3 by Treg is sufficient to drive Treg to a more effector-like phenotype, resulting in enhanced suppressive activity and increased tumor growth. These findings may help to reconcile previous reports that some Tim-3 antibodies enhance T cell responses *in vivo*, while expression of Tim-3 has a cell-intrinsic ability to enhance TCR signaling and T cell activation. Thus, we propose that Tim-3 regulates anti-tumor immunity at least in part through enhancement of Treg function. To our knowledge, this is the first example in which expression of a single co-stimulatory molecule is sufficient to drive differentiation of Treg in this manner.

## Introduction

Immune checkpoint blockade therapies like those targeting the PD-1/PD-L1 axis have resulted in dramatic gains in the treatment of some tumors, but in most settings only 20-30% of patients respond (Ferris et al., 2016; Hamid et al., 2013; Hargadon et al., 2018; Ribas and Wolchok, 2018). The mechanisms responsible for this resistance are still not clear. One barrier to successful immunotherapy can be the presence within the tumor microenvironment of significant numbers of regulatory T cells (Treg) (Tanaka and Sakaguchi, 2017). The glycoprotein <u>T</u> cell (or <u>t</u>ransmembrane) <u>i</u>mmunoglobulin and <u>m</u>ucin domain 3 (Tim-3; *HAVCR2*) has attracted significant attention as a potential negative regulator of T cell responses, including in tumors (Anderson, 2014; Du et al., 2017). Many groups have developed Tim-3 mAbs, with the goal of using these antibodies for immunotherapy of solid tumors, and early stage clinical trials are now underway. This effort has mainly focused on the role of Tim-3 in regulating CD8+ cytotoxic (CTL) T cells, although Tim-3 is also constitutively expressed by several other immune cell types, including mast cells, dendritic cells and some NK cells (Banerjee and Kane, 2018). At least four ligands have been described to bind to Tim-3: galectin-9, HMGB1, phosphatidylserine and CEACAM1, with virtually no studies directly comparing the effects of these different ligands. In addition, these ligands are also known to bind other receptors in addition to Tim-3.

Tim-3 is also expressed by a small subset (about 5%) of regulatory T cells (Treg) in the periphery (secondary lymphoid organs and peripheral blood), and interestingly the proportion of Tim-3^+^ Treg is significantly higher (4-60%) among tumor-infiltrating Treg (Gao et al., 2012; Jie et al., 2013; Liu et al., 2018b; Sakuishi et al., 2013; Shen et al., 2016). We also determined that these Tim3-expressing Treg, in contrast to paired Tim-3 negative Treg, express an array of genes normally associated with effector T cell function. Along with this effector Treg (eTreg) phenotype, Tim-3^+^ Treg also possess enhanced *in vitro* suppressive capacity. Since modulation of Tim-3 is being explored as a possible therapeutic strategy, it is important to understand the role of Tim-3 on this substantial population of Treg. Indeed Tim-3 appears to identify a particularly functional Treg subset and could act as a signaling mediator enforcing unique activity of the Treg expressing this protein (Gao et al., 2012; Gautron et al., 2014; Jie et al., 2013; Liu et al., 2018b; Sakuishi et al., 2013). Elimination or impairment of Tim-3^+^ Treg may represent an important mechanism by which mAb’s targeting Tim-3 enhance anti-tumor immunity (Liu et al., 2018a). In addition, expansion or activation of Treg may contribute to the phenomenon of hyper-progression, which occurs in a small subset of patients treated with checkpoint blockade (Kamada et al., 2019). However, it is still not known whether Tim-3 itself intrinsically modifies the phenotype or function of Treg on which it is expressed, or if its expression merely correlates with altered Treg phenotype and function.

Using a novel mouse model that allows us to drive Tim-3 expression in a Cre-dependent fashion, we have now found that expression of Tim-3 by Treg is sufficient to push Treg toward an effector Treg state, as revealed by surface protein expression and gene expression. Among the phenotypic changes we observe are enhanced gene and protein expression of IL-10 and a dramatic shift in expression of genes associated with metabolism, away from oxidative phosphorylation and toward glycolysis. At a functional level, Treg with enforced Tim-3 expression display enhanced *in vitro* suppressive activity. Even more striking, transplanted tumors grow more rapidly in mice with Treg-specific expression of Tim-3. Thus, Tim-3 possesses intrinsic activity that can reproduce a substantial proportion of the phenotype acquired by Tim-3+ tumor-infiltrating Treg.

## Materials and Methods

### Mice

C57BL/6 mice carrying a *Rosa26* knock-in of a flox-stop-flox cassette containing a Flag-tagged mTim-3 cDNA have been described (Avery et al., 2018). FoxP3-YFP-Cre mice were obtained from Dr. Dario Vignali; FoxP3-eGFP-Cre-ERT2 mice were obtained from The Jackson Laboratory. CD4-Cre transgenic mice on a C57BL/6 background were originally obtained from Taconic. C57BL/6 mice for breeding were obtained from Jackson Labs. For inducible Cre-mediated deletion with the Cre-ERT2 model, tamoxifen (MP Biomedical, LLC) was administered at 1 μg via the i.p. route on five successive days. Animals were used in accordance with University of Pittsburgh IACUC-approved protocols.

### Flow cytometry and antibodies

Single cell suspensions from organs and tumors were incubated with Live/Dead fixable stain (Tonbo Biosciences) and Fc Block (CD16/CD32, Tonbo, BioLegend) for 20 minutes, followed by staining for various cell surface markers. For analytic flow cytometry, cells were captured on a BD LSR-II and BD Fortessa. Data were exported and analyzed in FlowJo (versions 8 and 10). Cell sorting was carried out with a BD FACS Aria. Example of gating strategy used for flow cytometry analysis of spleen and lymph nodes is shown in **Fig. S1a**. Gating strategy used for tumor analysis is shown in **Fig. S1b**.

The following antibodies and reagents were used for flow cytometry: CD4 clone RM-4-5 (BD Biosciences, BioLegend, Tonbo), CD8α clone53-6.7 (BD, BioLegend, Tonbo), CD8β clone H35-17.2 (BD, BioLegend, Tonbo), CD19 Clone ID3 (BD, BioLegend, Tonbo), CD11b clone M1/70 (eBiosciences), CD11c clone HL3 (BioLegend), CD25 clone PC61 (BioLegend, Tonbo, BD), FoxP3 clone FJK16S (Invitrogen), CD44 clone IM7 (BD, Tonbo, BioLegend), CD62L clone MEL-14 (BioLegend, Tonbo), ICOS clone 398.4A (BioLegend), Ki67 clone B56 (BD), CTLA-4 clone UC10-4B9 (BioLegend), CD103 clone M290 (BD), Nrp-1 clone 3E12 (BioLegend), CD39 clone Duha 59, CD73 clone TY/11.8 (BioLegend), Tim-3 clone RMT 3-23 (BioLegend) and clone FAB1529 (R&D), Flag clone L5 (BioLegend), PD-1 clone RMP1-30 (BioLegend), IL-10 clone JES5-16E3 (BioLegend), CD90.2 clone 532.1 (BD, BioLegend), CD45.2 clone104 (BioLegend), Helios clone 22F6 (BioLegend), KLRG-1 clone 2F1 (BioLegend), Ly-6-G clone IA8 (Tonbo), GITR clone DTA-1 (eBiosciences), TIGIT clone 1G9 (BioLegend), LAG-3 clone C9B7W (Invitrogen), GARP clone F011-5 (BioLegend). CellTrace Violet was from Invitrogen.

Samples analyzed for transcription factors were first stained for Live/Dead staining along with Fc Block (CD16/32) followed by cell surface marker staining, fixed/permeabilized with the eBioscience FoxP3 staining kit (#00-5523-00), followed by intracellular protein staining.

### Transplantable tumor models

B16-F10 and MC38 tumor cells were obtained from D. Vignali and grown in RPMI1640 (for B16) or Dulbecco’s modified Eagle’s medium (DMEM; for MC38). Both types of media were supplemented with 10% FBS, penicillin, streptomycin, glutamine, pyruvate, HEPES and non-essential amino acids. Cell cultures were maintained at 37°C and 5% CO2. Mice (day six after tamoxifen/oil) were injected with B16F10 melanoma (1.25 × 10^5^ cells intradermally) or MC38 (5 × 10^5^ cells subcutaneously). Tumors were measured every three days starting at day five, in two dimensions, using digital calipers, and plotted as tumor area (mm^2^).

### Lymphocyte isolation from organs and tumors

Splenocytes and lymph node cells were isolated using two frosted glass slides for mechanical dissociation. Spleen cells were further processed for RBC lysis using Invitrogen RBC lysis buffer (eBioscience 00-4333-57), followed by filtering through 70 μM nylon mesh. Single cell suspensions of splenocytes and lymph node cells were then used for downstream experiments and staining for flow cytometry purposes. TIL from MC38 tumors were isolated by roughly chopping tumors, followed by incubation with collagenase D (Roche Diagnostics) and 0.2 mg/ml DNAse IV (Sigma-Aldrich) for 30 minutes at 37°C with constant gentle shaking. Digestion was quenched using complete DMEM followed by two washes with PBS. RBC lysis was performed, and cells were filtered through a 70 μM nylon mesh, before resuspension and incubation with Fc Block to prevent nonspecific binding.

### In vitro suppression assay

Tim-3 positive Treg were isolated by pooling spleen and LNs from tamoxifen-induced FSF-Tim-3 x Foxp3-eGFP-Cre-ERT2 mice; control Tim-3 negative Treg were isolated similarly from a tamoxifen-treated, Cre-only mice. Treg were sorted into Tim-3^+^ Treg (live CD4^+^YFP^+^CD25^Hi^Tim-3^+^) and control Tim-3^−^ Treg (live CD4^+^YFP^+^CD25^Hi^Tim-3^−^). Conventional T cells (Tconv; live CD3^+^CD8^+^) were sorted from a C57BL/6^Thy1.1^ mouse as responder cells and labeled with 5 μM CellTrace Violet (Invitrogen). Whole splenocytes from 6-8 week-old, wild-type C57BL/6 female mice were irradiated with 3,000 rads and used as antigen presenting cells (APCs). Responder cells and APCs were plated in triplicate at 5 × 10^4^ cells/well; co-cultures were set up with the following ratios of control Tim-3^−^ or Tim-3^+^ Treg: 1:2-1:16 Treg:Teff; 25,000-3,125 Treg (Polanczyk et al., 2005). T cell activation was achieved by adding anti-CD3 mAb (BioLegend) at a final concentration of 1 μg/ml and cells were co-cultured in 200 μl RPMI for 72 hrs. Stained cells were analyzed with a BD Fortessa flow cytometer and CellTrace Violet dilution was quantified using FlowJo. Suppression was calculated using the following formula, adapted from (Collison and Vignali, 2011): percent suppression = ((fraction of proliferated Tconv cells alone-fraction of proliferated Tconv cells with Treg)/(fraction of proliferated Tconv cells alone)*100).

### Gene expression analysis and RNAseq pipeline

Live CD4^+^CD25^+^GFP^+^ from WT mice, CD4^+^CD25^+^GFP^+^Tim-3^+^FLAG^+^ and CD4^+^CD25^+^GFP^−^Tim-3^+^FLAG^+^ cells from spleens of transgenic mice were sorted on a FACSAria directly into SmartSeq low-input RNA kit lysis buffer in a 96 well plate. DNA libraries were prepared using Nextera XT Kit and RNAseq was performed on an Illumina NextSeq 500 by The Children’s Hospital of Pittsburgh Sequencing core facility.

The reverse unstranded paired-end RNA-Seq reads of mouse regulatory T cells, generated by SMART-seq HT kit, were checked for presence of adapters and high-quality bases using FastQC (v 0.11.7). These high-quality reads were trimmed for adapters using Cutadapt (v 1.18). The trimmed reads were later mapped against the Ensembl mouse reference genome (GRCm38) using the HISAT2 (v 2.1.0) mapping tool. The output file from HISAT2 was converted from SAM format to BAM format using SAMtools (v 1.9). Counts for expressed genes were generated using HT-Seq (v 0.11.2) and were output in text format. These count text files were then imported into the Bioconductor R package, edgeR (v 3.24.1). The package was utilized to identify the differentially expressed genes based on the criteria of the genes having an expression count of absolute value log base 2 greater than 1 between two experimental conditions and a false discovery rate of less than 0.05. Based on this standard, between the 3 comparisons of WT Treg vs FoxP3^Hi^, WT Treg vs FoxP3^Lo^, and FoxP3^Hi^ vs FoxP3^Lo^ produced 5027, 5685, and 834 differentially expressed genes, respectively.

Analysis of single cell RNAseq data from human HNC TIL were carried out with an updated dataset based on that of Cillo et al. (Cillo et al., 2020).

### Statistical analysis

Experiments were analyzed using Student’s t-test for two-way comparisons. Statistical analyses were performed with GraphPad Prism. Tumor growth curves were analyzed using t-test for each time point and statistical significance was determined by the Holm-Sidak method, with alpha = 0.05. Each time point was analyzed individually, without assuming a consistent SD. P values are indicated as follows: \**P*◻<◻0.05, \*\**P*◻<◻0.01 and \*\*\**P*◻<◻0.005 and ****P<0.001 where statistical significance was found, and all data are represented as mean◻±◻s.d.

## Results

### Effects of enforced Tim-3 expression on the phenotype of circulating Treg

To probe the intrinsic effects of Tim-3 on Treg *in vivo*, we employed a mouse model that we had previously described(Avery et al., 2018) which carries a flox-stop-flox cassette followed by a full-length murine Tim-3 cDNA, knocked into the *Rosa26* locus (referred to as “FSF-Tim-3” mice). Upon expression of the Cre recombinase, cells carrying this allele express Tim-3 for the remaining life of that cell. Thus, we bred the FSF-Tim-3 to mice expressing a Cre/YFP fusion protein from a knock-in of the *Foxp3* locus (**Fig. 1a**) (Rubtsov et al., 2008). As expected, most (97-98%) FoxP3-expressing cells from the double transgenic mice constitutively expressed Tim-3, in contrast to the very low level of endogenous Tim-3 expression in Treg from the Cre-only control mice (**Fig. 1b**). Similar proportions of Tim-3^+^ Treg were observed in spleen and peripheral lymph nodes. We did note a small but statistically significant increase in the frequency and number of Treg in the FoxP3-Cre x FSF-Tim3 transgenic mice (**Fig. S2a**).

**Figure 1.**
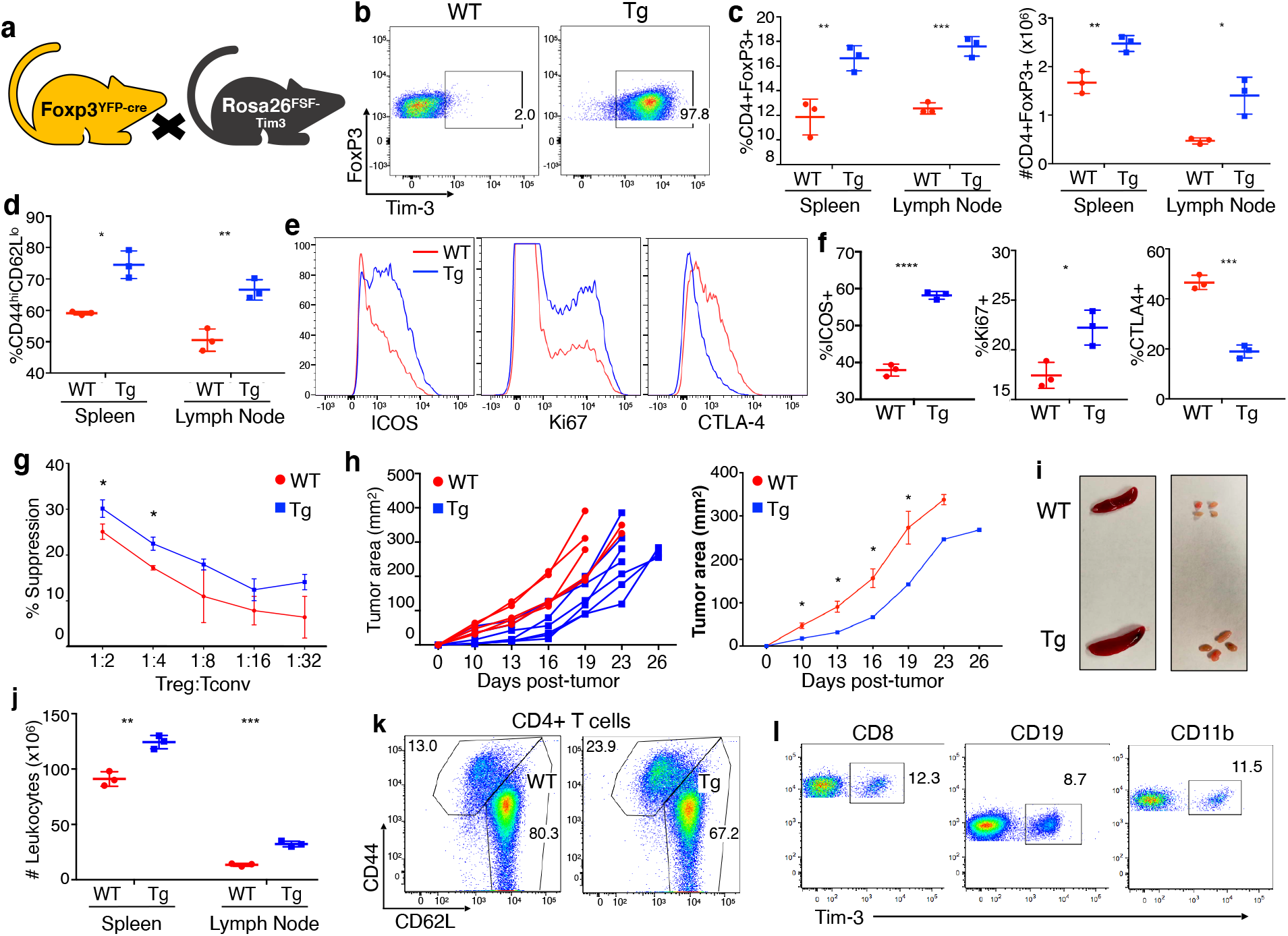
Phenotypic and functional effects of FoxP3-YFP-Cre-driven Tim-3 expression on Treg. (a) Crosses used to generate FoxP3-YFP-Cre x FSF-Tim-3 mice. (b) Efficiency of Tim-3 expression on Treg. Shown is a representative male mouse. Similar results were observed with Cre-homozygous female mice. (c) Ectopic Tim-3 expression via FoxP3-YFP-Cre resulted in an increase in the proportion of FoxP3^+^ cells among CD4^+^ T cells in spleen and lymph nodes. (d) Increase in activated Treg in the context of ectopic Tim-3 expression. (e-f) Expression of selected markers of Treg effector status and function in WT vs. Tim-3 Tg Treg. (g) *In vitro* suppression assay performed with FoxP3^−^YFP-Cre^+^ Treg sorted from Cre-only (WT) or Tim-3 Tg mice. (h) Growth of B16 tumors in WT vs. Tim-3 Treg Tg animals. Left – tumor growth in individual mice; right – average tumor growth. (i-j) Organ size and lymphocyte cell number of WT and Tim-3 Tg animals at eight weeks of age. (k) left – similar proportions of previously activated CD4^+^ T cells in spleen of WT vs. Tim-3 Tg animals. (l) Leaky expression of the Tim-3 transgene in multiple cell types from the spleen. Data are from a single experiment, representative of three experiments, with 3-4 mice per experiment, except for tumor experiments, which are representative of two experiments of 6-8 mice per experiment.

We next assessed the phenotype of Treg with or without induced Tim-3 expression. We first performed multi-color flow cytometry on T cells from spleen and peripheral lymph nodes of relatively young (eight weeks-old) mice expressing Cre, with or without the inducible Tim-3 allele. Thus, Tim-3 Tg Treg displayed higher baseline activation, based on expression of markers like CD44 and CD62L (**Fig. 1d; Fig. S2b**) and ICOS (**Fig. 1e-f**). Tim3-expressing Treg also appeared to be more proliferative, based on expression of Ki-67 (**Fig. 1e-f**). Curiously, however, constitutive expression of Tim-3 was associated with significant downregulation of CTLA-4 (**Fig. 1e-f**). Thus, overall, expression of Tim-3 was sufficient to drive a more-differentiated and activated phenotype in Treg residing in circulating Treg.

Acquisition of an effector Treg phenotype, as noted above, is associated with increased suppressive activity (Cretney et al., 2013; Teh et al., 2015). Thus, it was important to determine the functionality of Tim-3 Tg Treg. To do this, we first sorted Treg from control (FoxP3-YFP-Cre only) or Tim-3 Tg mice, based on expression of YFP-Cre. In parallel, we sorted WT naïve CD8^+^ T cells from the spleen of C57BL/6 mice. Proliferation was quantitated based on dye dilution of labeled CD8^+^ cells over the course of 72 hr co-culture with the WT or Tg Treg. Thus, we noted that Tim-3 Treg were capable of more potent suppression of WT effector T cells, compared with WT Treg **(Fig. 1g)**. Although *in vitro* suppression assays provide a useful reductionist snapshot of Treg activity, it is important to also assess Treg functionality *in vivo*. Past work implicated Tim-3^+^ Treg in helping to maintain a more suppressive tumor microenvironment (Gao et al., 2012; Jie et al., 2013; Liu et al., 2018b; Sakuishi et al., 2013). We therefore assessed the impact of enforced Tim-3 expression on growth of a transplantable tumor *in vivo*. Surprisingly, we found that growth kinetics of B16F10 melanoma tumors were slower in Tim-3 Treg transgenic mice, compared to WT control mice **(Fig. 1h)**. This finding is not consistent with the enhanced suppressive activity of Tim-3 Tg Treg in vitro, leading us to further explore the reasons for these divergent phenotypes. Upon further aging of Cre/FSF-Tim-3 double transgenic mice, we noted signs of autoinflammation, based on moderate splenomegaly and lymphadenopathy (**Fig. 1i-j**). However, this phenomenon did not appear to be due to an expansion of a particular cell type, as major cell populations were still present at similar proportions in the double transgenic vs. control animals **(Fig. S2c).**Inflammation in the transgenic mice was also associated with higher activation status of CD4 and CD8 compartment in both spleen and peripheral lymph nodes **(Fig. 1k; Fig. S2d).**

We considered whether lymphoid hyperplasia and reduced tumor growth might be linked to “leaky” expression of FoxP3-YFP-Cre outside of the Treg compartment, since it has been reported that such off-target expression may occur with this Cre strain (Franckaert et al., 2015). Indeed, when we directly examined different immune cell types, we saw widespread expression of the Tim-3 transgene (based on Tim-3 Flag staining) **(Fig. 1l)**. Thus, we observed expression of transgenic Tim-3 on significant numbers of both CD8^+^ (12%) and CD4^+^ non-Treg (40%) T cells, B cells (8.7%), and myeloid cells (11.5%). This suggests that the generalized lymphoid hyperplasia and baseline T cell activation in FoxP3-YFP-Cre transgenic mice is due to off-target Tim-3 expression. This appears to be unique to FoxP3-YFP-Cre, since we observed appropriate expression of transgenic Tim-3 in FSF-Tim-3 x CD4-Cre mice (Lee et al., 2001), i.e. on mature CD4^+^ and CD8^+^ T cells and Treg, but not B cells or myeloid cells (**Fig. S3a**). As expected, proportions of CD4^+^ and CD8^+^ T cells were normal in the periphery (**Fig. 2b**) and those T cells had unchanged numbers of previously activated cells (**Fig. 2c**). Nonetheless, we still observed a more activated phenotype and higher frequencies of Treg, relative to other lymphocytes, in both spleen and peripheral lymph nodes (**Fig. S3d**). As with FoxP3-YFP-Cre, CD4-Cre-dependent Tim-3 expression was associated with downregulation of CTLA-4 on Treg (data not shown). Taken together, these data suggested that transgenic expression of Tim-3 on circulating Treg was sufficient to drive these cells to a more effector Treg state, but the spurious expression in non-Treg did not allow us to rule out indirect effects due to effects on other cell types. Together with Fig. 1, these findings suggest that it is the spurious expression of transgenic Tim-3 in myeloid and/or B cells that drives the lymphadenopathy and splenomegaly.

**Figure 2.**
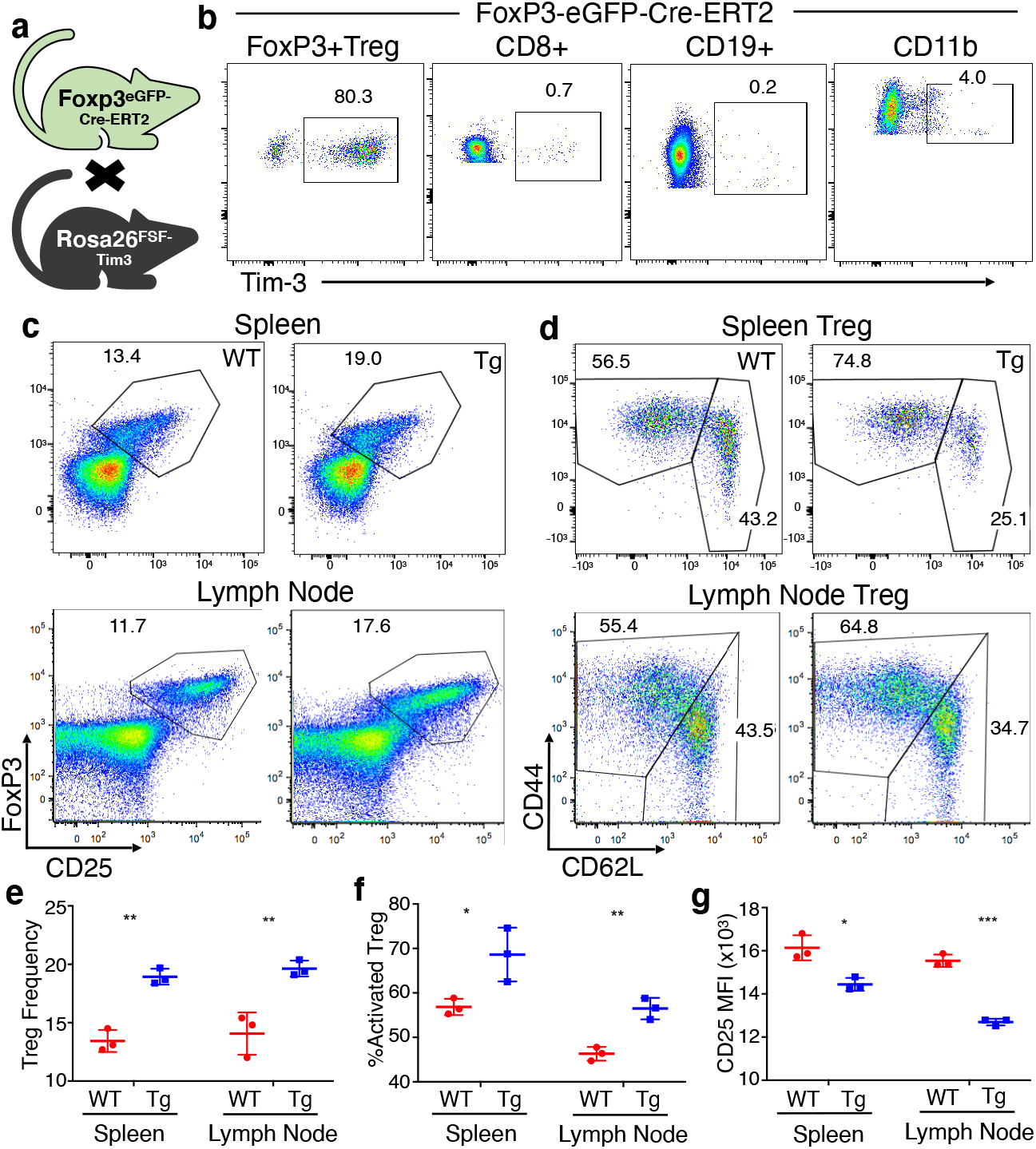
More-restricted expression and Treg activation phenotype of FoxP3-eGFP-Cre-ERT2 x FSF-Tim3 mice. (a) Crosses used to generate FoxP3-eGFP-Cre-ERT2 x FSF-Tim-3 mice. (b) Tim-3 transgene expression is restricted to CD4^+^ Treg when driven by the tamoxifen-inducible Cre. (c) Proportion of FoxP3^+^CD25^+^ Treg in spleen and LN of Tim-3 Tg vs. WT (Cre only) animals. (d) Activation status of Treg from the Tim-3 Tg vs. WT animals. (e) Quantitation of Treg frequency (among CD4^+^ T cells). (f) Quantitation of activated (CD44^+^CD62L^−^) Treg. (g) Levels of expression of CD25 (based on MFI of flow cytometry data). Mice were analyzed at 6-8 weeks of age. Representative of three experiments with 3-4 mice per experiment.

### Phenotypic effects of tamoxifen-inducible expression of Tim-3 on Treg

To drive restricted, but also temporally regulated, expression of Tim-3, we bred the FSF-Tim-3 mice to mice with a knock-in of a tamoxifen-inducible Cre (Cre-ERT2) fused to GFP in the *Foxp3* locus(Levine et al., 2014) (**Fig. 2a**). Indeed, when we crossed FSF-Tim3 mice to the FoxP3-eGFP-Cre-ERT2 mice, and administered tamoxifen, we did not observe any nonspecific induction of Tim-3 on non-Treg cells (**Fig. 2b**). Using the tamoxifen administration protocol described in *Materials and Methods*, females homozygous for the Cre and transgenic males had nearly 80 to 95% Tim-3^+^ Treg (**Fig. 2b**). In transgenic mice, there was a slightly higher frequency and number of FoxP3^+^ Treg in peripheral lymphoid organs at an early time point (day 7) (**Fig. 2c, e**), suggesting that enforced expression of Tim-3 on all FoxP3^+^ cells does not grossly impair the development or maintenance of Treg. Consistent with observations using the FoxP3-YFP model, expression of Tim-3 is sufficient to drive Treg towards a more effector-like phenotype. This is typified by the enhanced expression of both CD44 and ICOS (**Fig. 2d, f; Fig. 3a, Figs. S4-S5**), two known markers of effector Treg (eTreg) (Cretney et al., 2013; Smigiel et al., 2014). Induction of Tim-3 expression was also sufficient to alter other phenotypic aspects of Treg, with functional markers like CD103, Nrp-1, CD39 and CD73 showing increased expression (**Fig. 3, Figs. S4-S5**). By contrast, there were minimal or no differences in other functional markers like KLRG-1, Ki-67, GITR, GARP, TIGIT and Helios. Surprisingly, even though we observed apparent activation of Treg after induction of Tim-3, we also saw downregulation of CTLA-4 and CD25, along with a decrease in FoxP3 expression (**Fig. 2c, g; Fig. 3f, Figs. S4-S5**).

**Figure 3.**
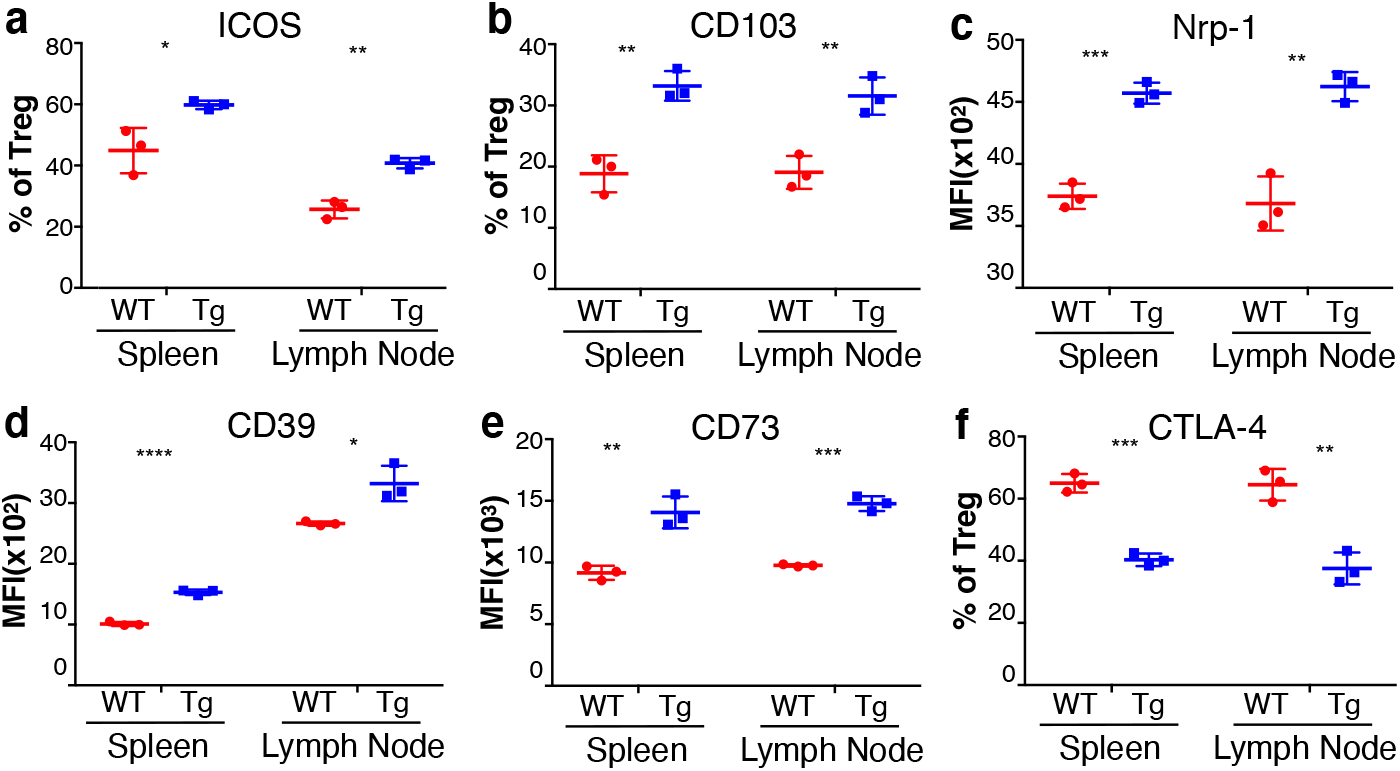
Detailed cell-surface phenotype of Treg from FoxP3-eGFP-Cre-ERT2 x FSF-Tim3 mice. (a-e) Statistically significant increases in ICOS, CD103, Nrp1, CD39 and CD73 were all observed on Tim-3 Tg Treg, in comparison to WT Treg (Cre-only). (f) By contrast, there was a corresponding decrease in expression of CTLA-4 on the Tim-3 Tg Treg. Mice were analyzed at 6-8 weeks of age. Representative of three experiments with 3-4 mice per experiment.

Based on results with the FoxP3-YFP-Cre model shown above, we wanted to determine if changes in the functional markers on Treg led to any gross phenotypic changes in the effector T cell compartment. Thus, at day seven after tamoxifen, there was no change in the frequency of CD4^+^ or CD8^+^ T cells in spleen or lymph nodes (**Fig. 4a, d**). Also, in contrast to the “leakier” FoxP3-YFP-Cre, the activation status (based on CD44 vs CD62L) of CD4^+^ (non-Treg) and CD8^+^ T cells in the spleen (**Fig. 4b, e**) and lymph node (**Fig. 4c, f**) was also unaffected. Finally, consistent with these phenotypic findings, the sizes of spleen and peripheral lymph nodes were also normal in the FSF-Tim-3 x FoxP3-eGFP-Cre-ERT2 mice (**Fig. 4g**). Thus, restricted and robust expression of Tim-3 on Treg in the periphery is sufficient to drive these cells to a more eTreg-like phenotype, without gross alterations in the activation status of conventional T cells.

**Figure 4.**
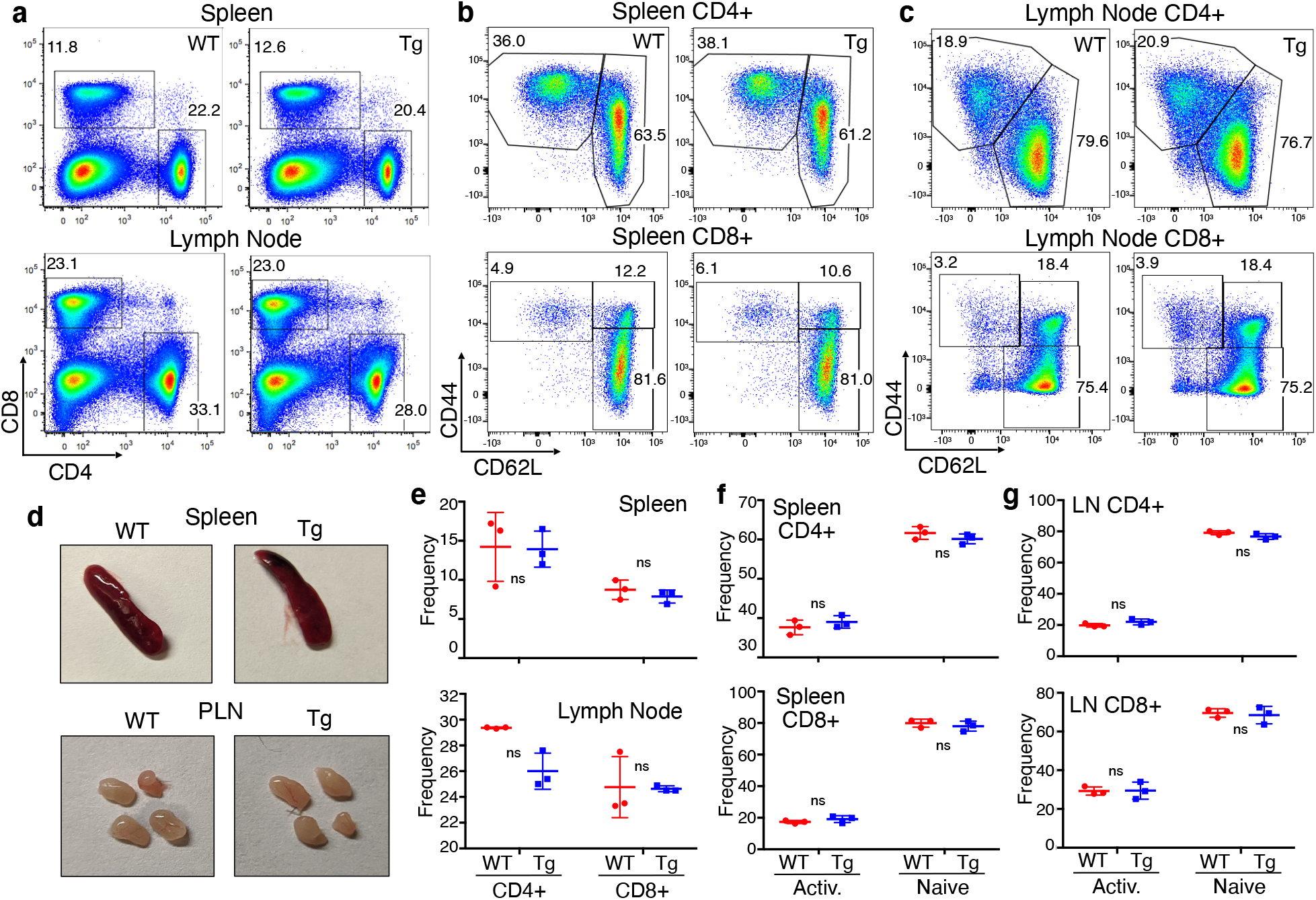
Phenotype of conventional T cells in peripheral lymphoid organs of FoxP3-eGFP-CRE-ERT2 x FSF-Tim-3 mice. (a) Proportion of CD4^+^ and CD8^+^ T cells in the spleen and lymph nodes of WT (FoxP3-eGFP-Cre-ERT2 only) and Tim-3 Tg animals, showing normal proportions of both cell types. (b-c) Proportions of antigen-experienced conventional T cells, based on CD44 and CD62L expression, again showing no evidence of basal lymphocyte activation in the Cre-ERT2 model. (d) Gross phenotypes of spleen and lymph nodes were also indistinguishable between WT and Tim-3 Tg animals. Quantitation of conventional T cell frequencies (e) and the proportions of antigen-experienced (CD44^+^) cells in spleen (f) and lymph node (g) from three animals per genotype. Representative of three experiments with 3-4 mice per experiment.

### Effects of Tim-3 on Treg homeostasis

Transient destabilization of Treg is thought to be important for licensing efficient effector T cell responses to pathogens, while aberrant or more profound Treg destabilization can lead to autoimmunity (Piconese and Barnaba, 2015; Sakaguchi et al., 2013; Shi and Chi, 2019). We therefore were interested in determining whether transgenic expression of Tim-3 altered Treg homeostasis *in vivo*. Experiments above were performed either with male mice or with female mice homozygous for the Cre transgene, as *Foxp3* is located on the X-chromosome and thus subject to random X-inactivation. In female mice carrying the FSF-Tim-3 allele and heterozygous for FoxP3-Cre, it is therefore expected that 50% of Treg in such mice will express transgenic Tim-3. Thus, after administration of tamoxifen to heterozygous females carrying FoxP3-eGFP-Cre-ERT2, we found that both Tim-3 induction and the GFP^+^ fraction in heterozygous transgenic mice were lower in number than heterozygous females carrying the Cre only, ranging from 20-35% of total Treg. This was seen both in unfixed cells, where we assessed FoxP3 indirectly based on the eGFP-Cre (**Fig. 5a-b**) and in fixed cells, where we stained for FoxP3 directly (**Fig. 5c-d**). We were therefore interested in whether the enforced expression of Tim-3 on Treg would alter their stability, a state usually associated with maintenance of high levels of FoxP3 expression, among other markers (Sakaguchi et al., 2013). We administered tamoxifen to WT (Cre-only) or Tim-3 Tg animals, then analyzed expression of FoxP3 and CD25 in CD4^+^ T cells from spleen and lymph nodes. By six weeks of age there was a statistically significant difference in the proportion of Treg in the spleen, but not in lymph nodes (**Fig. 5 e-f**). When we analyzed expression of FoxP3 and Tim-3 over time in Tg animals treated with tamoxifen, we noted a progressive decrease in FoxP3 expression, starting between three days and three weeks (**Fig. 5g**). By six weeks after tamoxifen treatment, very few FoxP3^hi^Tim-3+ cells remained in the spleen. Nonetheless, we did not observe any overt signs of the scurfy phenotype commonly associated with loss of Treg function (cachexia, runting, thickening/scaly skin). This suggests that either the remaining Tim-3^+^FoxP3^lo^ cells retain the ability to suppress autoimmunity or that such cells are replaced by new Treg coming from the thymus. To further explore if Tim-3 induction on Treg cells is sufficient to drive major changes in the basic characteristics of these cells, we performed bulk RNAseq of Treg cells from WT (FoxP3-eGFP-Cre-ERT2) and Tim-3^+^ transgenic (FoxP3-Cre x FSF-Tim3) Treg. In addition, we sorted both GFP^Hi^ (FoxP3^+^) and GFPl^Lo^ (FoxP3^−/lo^) cells and both populations were compared to WT Treg (**Fig 5h**). The latter population was included to assess the gene expression profile of the apparently destabilized Treg discussed above.

**Figure 5.**
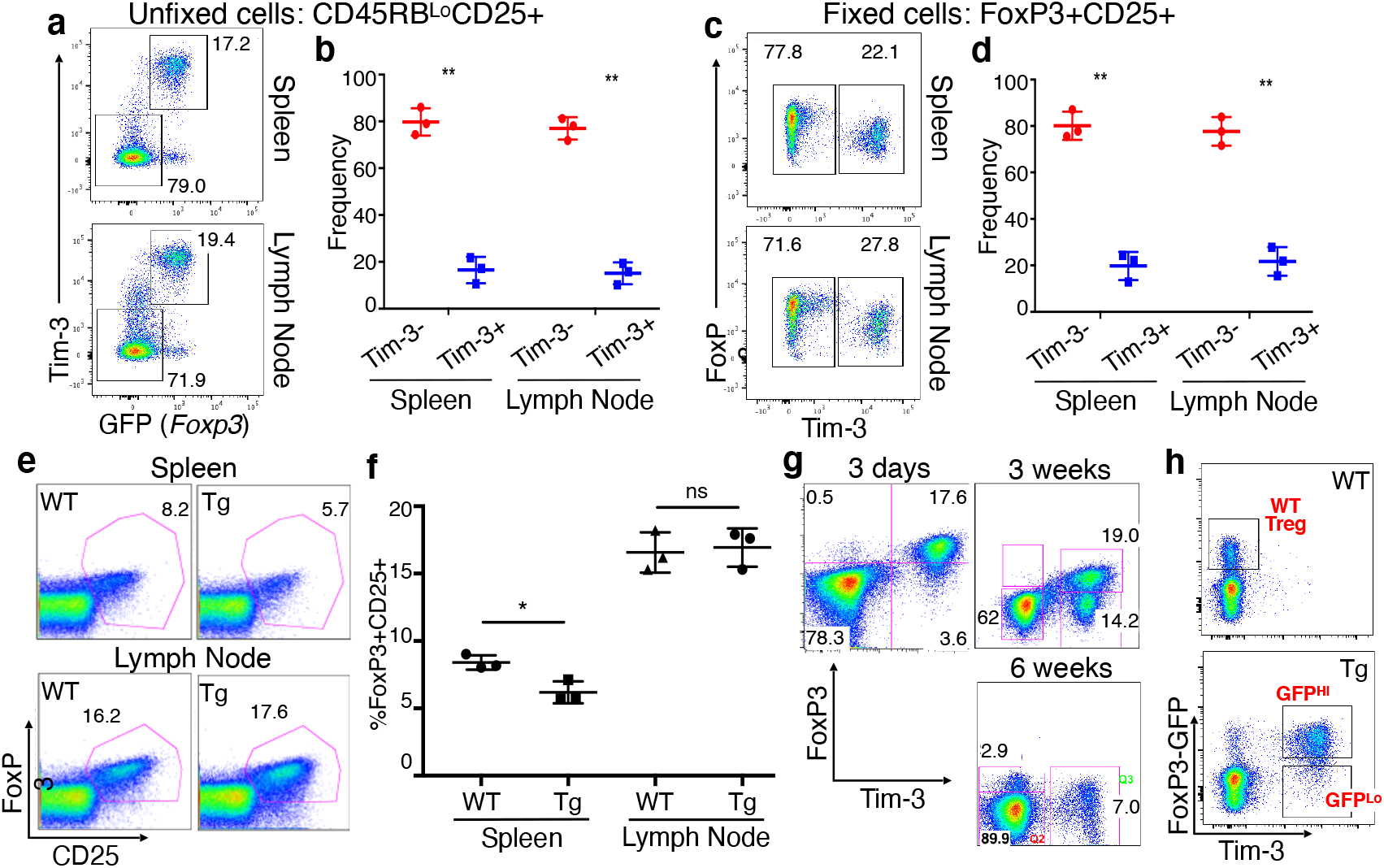
Evidence of partial Treg destabilization in the spleen after Tim-3 induction. (a-b) Expression of the Tim-3 transgene and GFP (as a surrogate for FoxP3) in unfixed Treg from FoxP3-eGFP-Cre-ERT2 heterozygous female mice. (c-d) Expression of the Tim-3 transgene and FoxP3 protein in fixed Treg. (e-f) Proportion of FoxP3^+^CD25^+^ Treg among CD4^+^ T cells in spleen and lymph node, showing a slight but statistically significant decrease in the proportion of splenic, but not lymph node, Treg in the Tg animals, by six weeks of age after tamoxifen administration. (g) FoxP3 and Tim-3 expression among splenic CD4^+^ T cells at the indicated times after tamoxifen administration. Expression of Tim-3 is permanent once Cre-mediated recombination occurs at the modified *Rosa26* locus, making Tim-3 expression a faithful fate-tracking reporter. (h) FoxP3-eGFP-Cre and Tim-3 expression in WT (Cre-only) vs. Tim-3 Tg Treg, indicating the three cell populations sorted for RNA sequencing.

### Effects of Tim-3 expression on Treg gene expression

After sorting the cell populations indicated in **Fig. 5h**, RNA was made from the sorted cells and prepared for RNA sequencing. We found that Tim-3 induction led to major changes in the Treg cells transcript signature, even in the Tim3-expressing Treg that still expressed high levels of FoxP3, as illustrated in the volcano plot in **Fig. 6a**. Based on Kyoto Encyclopedia of Genes and Genomes (KEGG) analysis of the top 500 differentially expressed genes in the Tim-3+FoxP3^hi^ population, major housekeeping processes in the Treg seem to be altered due to expression of Tim-3, including translation (i.e. ribosomal genes) and metabolism (**Fig. 6b, Figs. S6-S7**). Relative expression of genes of interest from these modules, as well as cytokines/receptors and transcription factors, are shown in **Fig. 6c**, where we also compared their expression in the Tim-3+FoxP3^Lo^ cells (see also **Figs. S6-S7**). Of note, the message encoding the suppressive cytokine IL-10 is massively upregulated in the Tim-3 Tg Treg (see also the volcano plot in **Fig. 6a**), and there were broad decreases in ribosomal and oxidative phosphorylation/mitochondrial genes, with a corresponding increase in genes involved in glycolysis. Consistent with the results discussed in **Fig. 5**, the GFP^Lo^ (i.e. FoxP3^Lo^) but not GFP^Hi^ Tim-3^+^ Treg had greatly reduced *Il2ra* message, which encodes CD25. Finally, consistent with the acquisition of eTreg status by Tim-3 Tg Treg, we observed increased expression of the transcription factors Eomes, Myc and T-bet (*Tbx21*).

**Figure 6.**
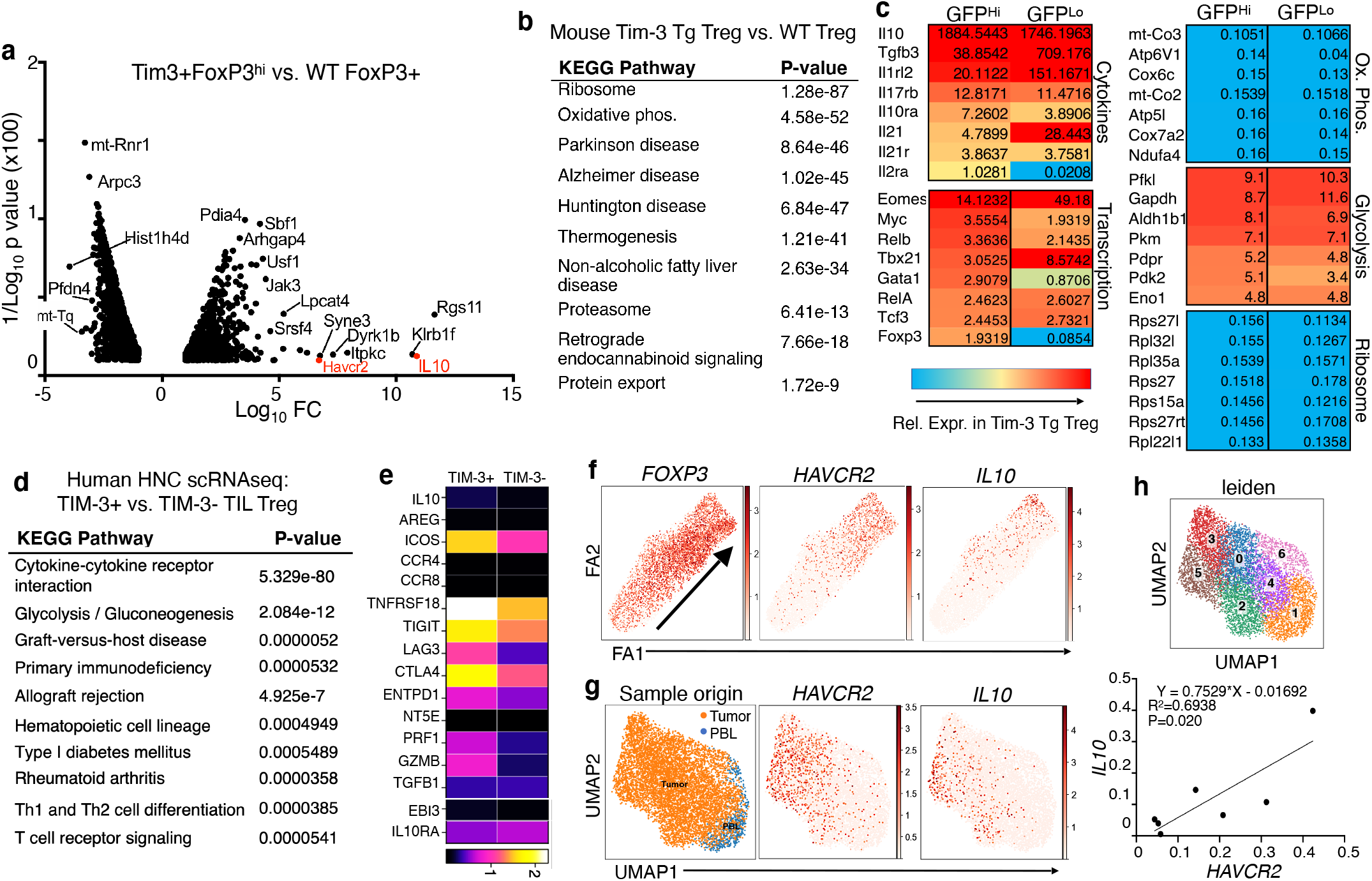
Effects of enforced Tim-3 expression on Treg gene expression. (a) Volcano plot of RNA sequencing data comparing Tim-3^+^FoxP3^hi^ cells from Tg animals to FoxP3^+^ cells from WT (Cre-only) animals. Only comparisons with a p value <0.05 are shown. (b) KEGG pathway analysis of the top 500 differentially expressed genes by fold change and FDR (either up or down), comparing FoxP3^hi^Tim-3^+^ cells to WT FoxP3^+^ cells. (c) Differential expression of selected genes in the indicated pathways, comparing either Tim-3^+^GFP^hi^ (i.e. FoxP3^hi^) or Tim-3^+^GFP^lo^ cells to WT Treg from FoxP3-Cre-GFP mice. Mice were injected with tamoxifen at 6-8 weeks of age and analyzed between three and six weeks of age. (d) KEGG analysis of human HNC TIM-3^+^ vs. TIM-3^−^ TIL Treg, based on scRNAseq, using the top 100 upregulated and top 100 downregulated genes. (e) Heat map showing relative expression of selected genes in TIM-3^+^ vs. TIM-3^−^ human Treg from the scRNAseq dataset. (f) Pseudotime analysis of TIL Treg from HNC patient scRNAseq data, showing that increased *HAVCR2* and *IL10* expression are associated with differentiation (arrow trajectory) of Treg. (g) UMAP analysis of human HNC patient Treg for *HAVCR2* and *IL10*. (h) top – definition of clusters from UMAP analysis shown in panel g; bottom – regression analysis of *HAVCR2* vs. *IL10* expression in the various Treg clusters.

We previously reported that about 50% of human tumor-infiltrating Treg express TIM-3 and that these cells possess enhanced suppressive function, compared to matched Treg not expressing TIM-3 (Jie et al., 2013; Liu et al., 2018b). Thus, we wanted to determine if the pathways identified as driven by Tim-3 expression by RNAseq analysis of our Tg mouse model are also observed in human TIL Treg. Recently, we published a comprehensive study of leukocytes isolated from HNC patient blood and tumors (Cillo et al., 2020). Analysis of an updated scRNAseq dataset, using the top 100 differentially expressed genes (**Fig. S8**), revealed overlap with the mouse Tim-3 Tg Treg analysis. Of note, two of the most significantly altered pathways, by KEGG analysis, were the ribosome pathway and glycolysis (**Fig. 6d**). Looking at the relative expression of individual genes associated with effector Treg, we noted upregulation of messages encoding IL-10 and ICOS, among other genes (**Fig. 6e**). One divergence from the mouse data is the upregulation of CTLA-4 in TIM-3^+^ human TIL Treg, which contrasts with the lower expression of Ctla-4 in Tim-3+ Tg mouse Treg. Given the prominence of IL-10 upregulation (both message and protein) in mouse Tim-3 Tg Treg, we examined the expression of IL-10 in human TIL Treg in more detail. We also wanted to assess whether TIM-3^+^ TIL Treg represent a more-differentiated state, as suggested by the analysis of Tim-3 Tg Treg over time in the mouse model. Thus, we performed pseudotime analysis of FOXP3-expressing cells from the scRNAseq dataset. As shown in **Fig. 6f**, pseudotime progression of TIL Treg (indicated by the diagonal arrow) is associated with increased expression of both *HAVCR2* (the gene encoding TIM-3) and IL10. Consistent with the enrichment of Treg in tumors, most of the Treg in our dataset were of tumor origin, with a minor fraction from the blood (**Fig. 6g**). Overlaying the expression of Tim-3 onto the Treg UMAP, we again observed a significant number of tumor-infiltrating, but not PBL, Treg expressing *HAVCR2* (**Fig. 6g**), which again appeared to correlate with expression of *IL10*. This same UMAP analysis revealed the presence of seven distinct subsets of Treg in the tumors (**Fig. 6h**, top). When we performed linear regression analysis, we found a significant correlation between *HAVCR2* and *IL10* expression within these clusters (**Fig. 6h**, bottom).

### Expression of Tim-3 drives enhanced Treg function

In addition to the phenotypic characterization of Tim-3 Tg Treg, it was important to also assess the functional effects of Tim-3 expression on Treg. We first tested the *in vitro* suppressive capacity of Tim-3 positive vs. Tim-3 negative Treg cells, using a standard *in vitro* suppression assay. FoxP3^+^ cells were sorted (based on GFP expression) were sorted from FoxP3-eGFP-Cre-ERT2 mice, with or without the FSF-Tim3 transgene. Thus, consistent with results described above using FoxP3-YFP-Cre (**Fig. 1**), Treg from the Tim-3 transgenic mice were more active in this assay, i.e. co-culture with the Tim-3 Tg Treg suppressed the proliferation of conventional CD4 and CD8 cells to a greater extent than WT Treg (**Fig. 7a-b**). To further define the functional competence of Treg with enforced Tim-3 expression, we implanted tumors into control (FoxP3-eGFP-Cre-ERT2) or Tim-3 Tg mice. We chose two tumor models for their different kinetics and relative immunogenicity – B16 melanoma, which is relatively aggressive; and MC38 carcinoma, which grows somewhat more slowly and is more immunogenic and responsive to immune checkpoint blockade (Woo et al., 2012). Thus, B16 tumors grew with typical kinetics in WT (FoxP3-eGFP-Cre-ERT2 only) mice (**Fig. 7c**), and this was not affected by administration of oil control (the solvent in which tamoxifen is dissolved). By contrast, even this aggressive tumor grew more quickly in Tim-3 Tg animals that were given tamoxifen to induce Cre-mediated recombination, and Tim-3 expression, in Treg (**Fig. 7c**). We observed an even more dramatic effect of Tim-3 expression on Treg in the MC38 tumor model (**Fig. 7d**). These effects of Treg-specific Tim-3 expression *in vivo* correspond well with the enhanced effector Treg phenotype and suppressive function.

**Figure 7.**
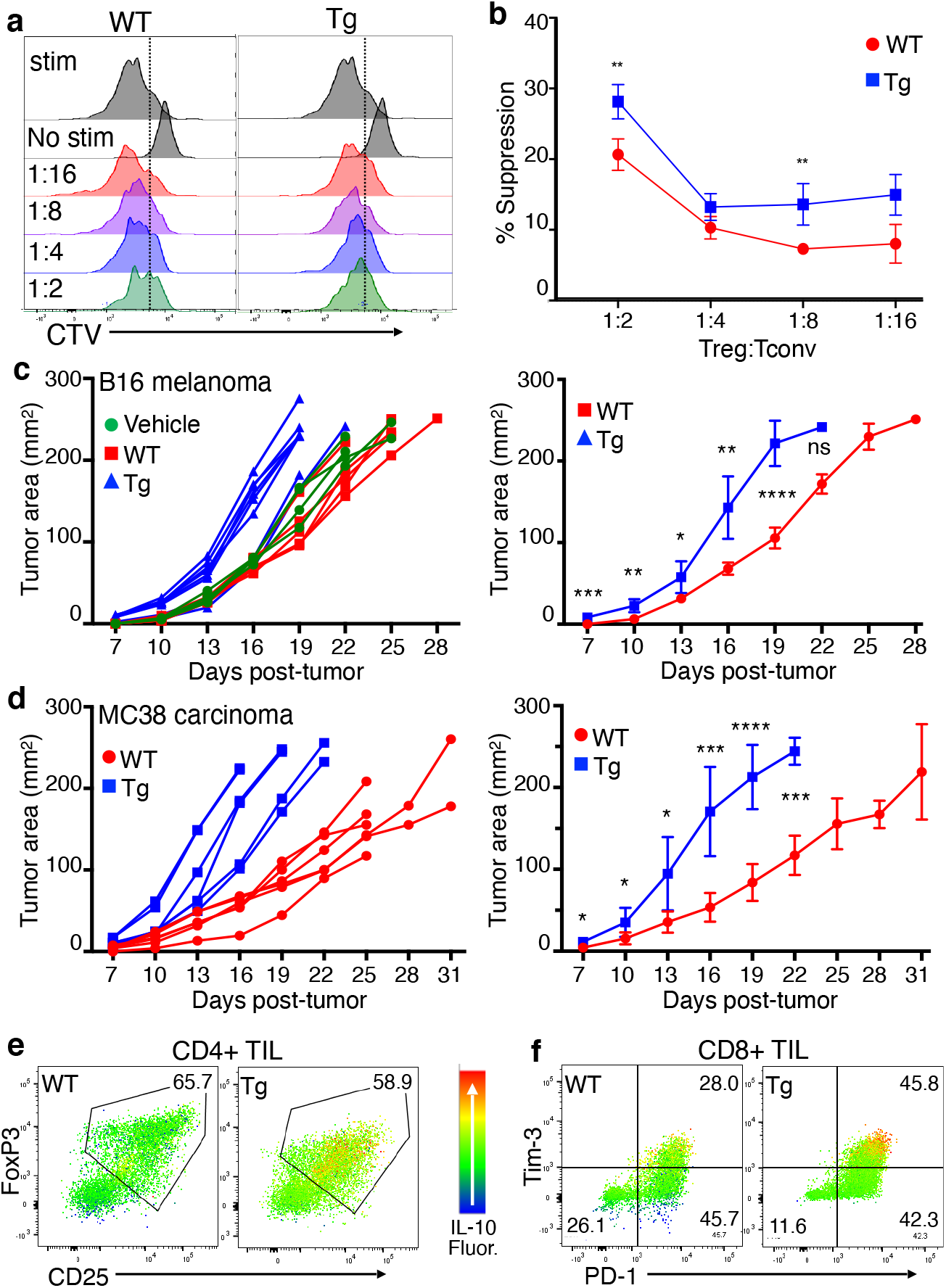
Treg with enforced Tim-3 expression display enhanced suppressive activity and IL-10 production. (a-b) GFP^+^ cells were sorted from Tim-3 Tg or WT (Cre-only) mice and mixed with sorted naïve conventional T cells from BL/6 mice for an *in vitro* suppression assay. Panel A shows representative CellTrace Violet (CTV) dilution, used to quantitate proliferation of the target conventional T cells. Panel B shows average suppression of triplicate samples over a range of ratios of Treg to conventional T cells. (c-d) Effects of Treg Tim-3 induction on transplanted tumor growth. The indicated mice were treated with tamoxifen or vehicle, followed by injection of B16 (c) or MC38 (d) tumors. Growth curves for individual animals are shown (left panels), as well as average tumor growth (right panels). (e-f) Effects of Treg Tim-3 expression on IL-10 production by exhausted CD8^+^ conventional T cells (e) or CD4^+^ Treg (f). Relative IL-10 expression heat-mapped onto Tim-3 x PD-1 expression for exhausted T cells or FoxP3 x CD25 expression for Treg. Representative of two experiments of four mice each, with six-week-old mice.

Treg employ multiple mechanisms to effect suppression of conventional T cell function. One of the most critical factors in this regard, including in the tumor microenvironment, is the regulatory cytokine IL-10 (Chen et al., 2016; Yano et al., 2019). Previous studies suggested that expression of Tim-3 is linked to increased IL-10 expression on both effector and regulatory T cells, with IL-10 expression by CD8^+^ T cells also being associated with T cell exhaustion (Gorman and Colgan, 2018; Gupta et al., 2012; Jin et al., 2010; Sakuishi et al., 2013; Zhu et al., 2015). We therefore examined the expression of IL-10 in CD8^+^ and FoxP3^+^ TIL isolated from MC38 tumors implanted into control or Tim-3 Tg mice. When we plotted the expression of IL-10 heat-mapped onto flow cytometry plots for Tim-3 and PD-1, which identify the most-exhausted CD8^+^ T cells in the tumor microenvironment, we found that Tim-3^+^PD-1^+^ effector T cells were enriched for IL-10 expression when Tim-3 was induced on Treg (**Fig. 7e**). In addition, there was a higher proportion of these PD-1^+^Tim-3^+^CD8^+^ T cells in tumors implanted into the Treg Tim-3 Tg animals. We also examined IL-10 expression in Treg from these animals. As shown in **Fig. 7f**, Treg (FoxP3^+^CD25^+^) from Tim-3 Tg animals showed higher expression of IL-10, compared with Cre-only control animals. Thus, constitutive expression of Tim-3 on Treg not only drives the Treg themselves to produce more IL-10, but it also leads to increased exhaustion and IL-10 production amongst CD8^+^ TIL T cells.

## Discussion

Tim-3 has attracted significant attention as a possible target for immune checkpoint blockade therapy, based on its co-expression, with PD-1, on a subset of exhausted T cells. Some pre-clinical evidence suggests that simultaneous targeting of these two proteins may enhance the sub-optimal responses to PD-1 blockade seen in most patients. However, Tim-3 is highly expressed by about 50% of Treg in the TME, and this expression is associated with more potent Treg suppressive activity. Here we set out to address whether Tim-3 itself can drive phenotypic and functional changes in Treg, including whether expression of Tim-3 makes Treg more potent regulators of effector T cells and the anti-tumor immune response. We found that transgenic expression of Tim-3 on Treg is indeed sufficient to make circulating Treg acquire a cell-surface phenotype reminiscent of effector Treg (eTreg). In addition, these Tim3-expressing peripheral Treg are more suppressive, a phenotype which also results in enhanced tumor growth. Consistent with previous studies that characterized eTreg as a more-differentiated and highly suppressive subset (Cretney et al., 2013; Teh et al., 2015), Tim-3 Tg Treg displayed higher expression of cell-surface protein like ICOS and CD44 and decreased CD62L. We also noted significantly higher expression of proteins known to help mediate Treg function, including the ecto-nucleases CD39 and CD73.

One of the more-unexpected effects of Tim-3 expression on Treg was the down-regulation of CTLA-4, which occurred relatively quickly after induction of Tim-3 by tamoxifen, especially since this occurred in the context of enhanced Treg suppression. Our RNA sequencing analysis revealed that this is due at least in part to a decrease in *Ctla4* message. Downregulation of CTLA-4 was also accompanied by corresponding decreases in CD25 and even FoxP3, which regulates expression of both CTLA-4 and CD25. This is precedent for this type of dichotomy between CTLA-4 expression and Treg suppression, as inducible knockout of CTLA-4 still allowed for normal Treg suppressive activity, both *in vitro* and *in vivo* (Paterson et al., 2015). Interestingly, this split phenotype was attributed in part to an upregulation of a number of genes, including IL-10 and ICOS (Paterson et al., 2015), both of which we also see upregulated in our Tim-3 Tg Treg model. Along with an enhanced effector phenotype, we noted an apparent de-stabilization of Treg, specifically in the spleen, based on expression of FoxP3 and CD25. Intriguingly, the shift from resting Treg to ICOS^+^ eTreg has been shown to correlate with destabilization, based on a sophisticated lineage tracing approach (Zhang et al., 2017). This shift was also shown to be associated with upregulation of T-bet, a transcription factor that regulates effector function of conventional T cells and Treg. Of note, Tbet message (encoded by *Tbx21*) was upregulated in Treg constitutively expressing Tim-3.

Most of the above-described changes in Treg biology were seen upon Cre-dependent expression of Tim-3 driven by either the older FoxP3-YFP-Cre model or the tamoxifen-inducible FoxP3-eGFP-Cre-ERT2 model; similar phenotypic changes were also observed with CD4-Cre. One exception was the difference in tumor growth, with the YFP-Cre model resulting in slower tumor growth, while the tamoxifen-inducible eGFP-Cre model yielded faster tumor growth. Based on the leakiness in Cre-mediated recombination that we observed, which was consistent with previous reports (Franckaert et al., 2015), we conclude that spurious expression of Tim-3 on non-Treg, including effector T cells, B cells and myeloid cells, exerted a dominant pro-inflammatory effect that predominated over the anti-inflammatory effect via enhanced Treg function. We were not able to localize the lymphoid hyperplasia to an expansion of any one cell type, so the proximal cause of this phenomenon remains unclear. However, previous work from our group and others (Anderson et al., 2007; Avery et al., 2018; Jayaraman et al., 2010; Phong et al., 2015; Qiu et al., 2012; Sada-Ovalle et al., 2012) provides compelling evidence that Tim-3 can increase the activation and/or effector function of multiple immune cell types, including T cells, DC’s, macrophages and mast cells.

How do the effects of Tim-3 on Treg biology fit within the wider context of Tim-3 function? As described above, while there are indications from multiple studies that Tim-3 can enhance immune cell function, other studies support a model of Tim-3 as a negative regulator, especially of exhausted T cells in the contexts of chronic viral infection or within the tumor microenvironment (Anderson et al., 2016; Banerjee and Kane, 2018; Du et al., 2017). We have previously discussed the possibility that some mAb’s to Tim-3 may have agonistic or agonistic function, especially in light of the fact that most studies have not explicitly addressed the role of blocking specific ligands in the function of these antibodies (Banerjee and Kane, 2018; Ferris et al., 2014). The ability of Tim-3 to enhance the suppressive activity of Treg suggests another mechanism by which Tim-3 mAb blocking activity may enhance immune responses. Thus, in the tumor context, such antibodies could interfere with the ability of Tim-3 to drive increased Treg suppressive activity, indirectly promoting the ability of CD8^+^ T cells to drive anti-tumor immunity. How these effects are mediated at the level of Tim-3 signaling is less clear at this point. The inhibitory signaling of Tim-3 has been postulated to proceed through *in trans* interactions or signaling through various molecules, including Bat3, CD45, CEACAM-1 and a long-noncoding RNA (Clayton et al., 2014; Huang et al., 2014; Ji et al., 2018; Rangachari et al., 2012). However, the increased accumulation of eTreg, as read-out by levels of cell-surface markers like CD44 and CD62L, as well as the shift toward a more glycolytic phenotype, all suggest involvement of a more active signaling process. We suggest that this is due at least in part to enhanced activation of Akt/mTOR signaling, which we previously reported as being increased in T cell lines and primary effector T cells expressing Tim-3 (Avery et al., 2018; Lee et al., 2011). Other groups have reported evidence for a co-stimulatory role for Tim-3 in T cells, including upregulation of mTOR activation (Gorman and Colgan, 2018; Gorman et al., 2014; Sabins et al., 2017). Interestingly, in a gene expression analysis of WT vs. Tim-3 KO CD4^+^ T cells, one of the most highly upregulated genes was IL-10 (Gorman and Colgan, 2018), so this may be a widespread target of Tim-3 signaling.

Further complicating the deconvolution of the *in vivo* function of Tim-3, including within the tumor microenvironment (TME), is the apparent existence of multiple ligands for the ecto domain of this protein. These various ligands (galectin-9, HMGB1, PS and CEACAM1) may intrinsically impact Tim3-dependent signaling and cellular function in different ways and in a context-dependent manner, e.g. in the TME vs. other peripheral tissues. The effects may then be propagated to other cell types in the environment, again in a context-dependent manner. Thus, we observed a correlation between expression of Tim-3 and IL-10 in Treg. We also noted an increase in IL-10 production by exhausted effector CD8^+^ T cells in that context, so the Tim-3^+^ Treg likely exert suppressive effects on other cell types in the TME. For example, the decreased level of CTLA-4 on Tim-3^+^ Treg may promote T cell over-stimulation, leading to exhaustion, due to more sustained expression of B7 family members in the TME (Greenwald et al., 2005; Zang and Allison, 2007). A dramatic upregulation of Tim-3 expression by Treg in the TME, compared with secondary lymphoid organs, has been a consistent finding across multiple studies (Jie et al., 2013; Liu et al., 2018b; Sakuishi et al., 2013; Shen et al., 2016). While our study addresses the function of Tim-3 on Treg, it does not answer the question of *how* these cells acquire expression of Tim-3. There are at least two mechanisms by which this might occur. One is the recruitment and expansion of circulating or tissue Tim-3^+^ Treg to tumors. We do observe a small fraction of Tim-3^+^ Treg in spleen and lymph nodes of normal mice, and also within the peripheral blood of human cancer patients (Liu et al., 2018b). We have not examined expression of Tim-3 on Treg from non-lymphoid tissue (e.g. from skin). However, there is published evidence for increased expression of immune checkpoint molecules on tissue-resident memory T cells (Gamradt et al., 2019; Weisberg et al., 2019). What about *de novo* acquisition of Tim-3 expression by TIL Treg? Although factors that regulate Tim-3 expression have not been exhaustively defined, several cytokines and transcription factors have been shown to promote Tim-3 expression on T cells. The transcription factors T-bet and NFIL3 can apparently help drive expression of Tim-3 on effector T cells in response to IFN-γ and IL-27, respectively (Anderson et al., 2010; Zhu et al., 2015). Additional factors likely remain to be revealed.

Two of the pathways that are most closely linked to Treg-specific expression of Tim-3 are cellular metabolism and protein translation. There is now an extensive literature demonstrating the roles of shifts in metabolic pathways among different subsets of T cells and during transitions between different activation and effector states of these same cells. For example, Treg appear to rely predominantly on mitochondrial metabolism, including lipid oxidation, while actively suppressing glycolysis, in a FoxP3-dependent manner (Shi and Chi, 2019). Thus, the upregulation of glycolysis genes, along with downregulation of oxidative phosphorylation genes, in Tim3-expressing Treg may be due at least in part to the downregulation of FoxP3. However, there could also be more direct effects of Tim-3 expression, since we previously reported that ectopic expression of Tim-3 in T cells can enhance activation of the mTOR pathway (Avery et al., 2018; Lee et al., 2011), which is known to promote glycolysis in T cells (Pollizzi and Powell, 2014). There is a paucity of literature regarding the regulation of protein translation in T cells, and especially in Treg, so the relevance of the Tim-3 effects on translation is less clear. Nonetheless, a recent report documented a role for ribosome biogenesis in controlling Treg activation and function, as Treg-specific knockout of Noc4L led to a scurfy-like phenotype and lower expression of ICOS and CTLA-4 (Zhu et al., 2019).

Finally, what is the relevance of Tim-3^+^ Treg in the context of tumor immunotherapy? We and others previously reported that TIL Treg upregulate not only Tim-3, but also PD-1. Tim-3 is an emerging target for tumor immunotherapy and PD-1 is now a first-line therapy for multiple tumor types, although overall response rates to checkpoint blockade therapy are still only on the order of 25% for single-agent regimens. Given the highly suppressive nature of Tim-3^+^PD-1^+^ Treg, therapies that specifically target these cells in the tumor microenvironment could be particularly effective. In addition, it has been suggested that there is a role for PD-1^+^ TIL Treg in the phenomenon of “hyper-progression,” which can sometimes occur in response to immunotherapy (Kamada et al., 2019). Thus, care should be taken to avoid Tim3-specific interventions that enhance the function of tumor Treg. Conversely, tumor-specific inhibition of this pathway may provide a target for selective enhancement of anti-tumor immunity, without attendant autoimmunity. This leads us to propose a unified model for Tim-3 function, as a co-stimulatory molecule with the ability to enhance the function of multiple immune cell types, including effector T cells, Treg, mast cells and some myeloid cells. The net effects of modulating Tim-3 function *in vivo* will therefore depend upon the relative expression of Tim-3 on these cell types, their functional states and access to Tim-3 ligands and modulating agents like monoclonal antibodies.

## Acknowledgements

We thank: Shunsuke Kataoka and Kyle McGraw for helpful discussions; Mia Simms for performing mouse genotyping; Lawrence Andrews, Angela Gocher and Elisa Ruffo for technical assistance with tumor experiments and Treg suppression assays; Creg Workman for input into experimental design; The Dept. of Immunology flow facility, especially Dwayne Falkner, for assistance with cell sorting; Health Sciences Sequencing Core at UPMC Children’s Hospital of Pittsburgh; Tim Hand, Mark Shlomchik and Dario Vignali for providing antibodies; Reinhardt Hinterleitner and Marlies Meisel for advice on intracellular IL-10 staining. Support from NIH/NCI award CA206517 (RLF and LPK); BM is supported by NCI training grant T32CA082084 (PI-Vignali); other support from NCI SPORE CA097190.

## Author Contributions

HB designed studies, performed most experiments and drafted and edited the manuscript. HN-R performed Treg suppression assays. AK performed analysis of human scRNAseq data and provided input on data presentation. BM assisted with design and analysis of tumor experiments. AC and URC performed analysis of bulk RNAseq from mouse Treg. ALS-W provided technical assistance and edited the manuscript. LV and RLF provided critical input into analysis of human scRNAseq data. LPK obtained funding for the project, conceived of initial study design and edited the manuscript.

## Competing Interests

The authors declare that they have no relevant conflicts of interest to report.

## Notes

### Competing Interest Statement

The authors have declared no competing interest.

