## Supplemental Figures for "Expression of Tim-3 drives naïve Treg to an effector-like state with enhanced suppressive activity"

Figure S1

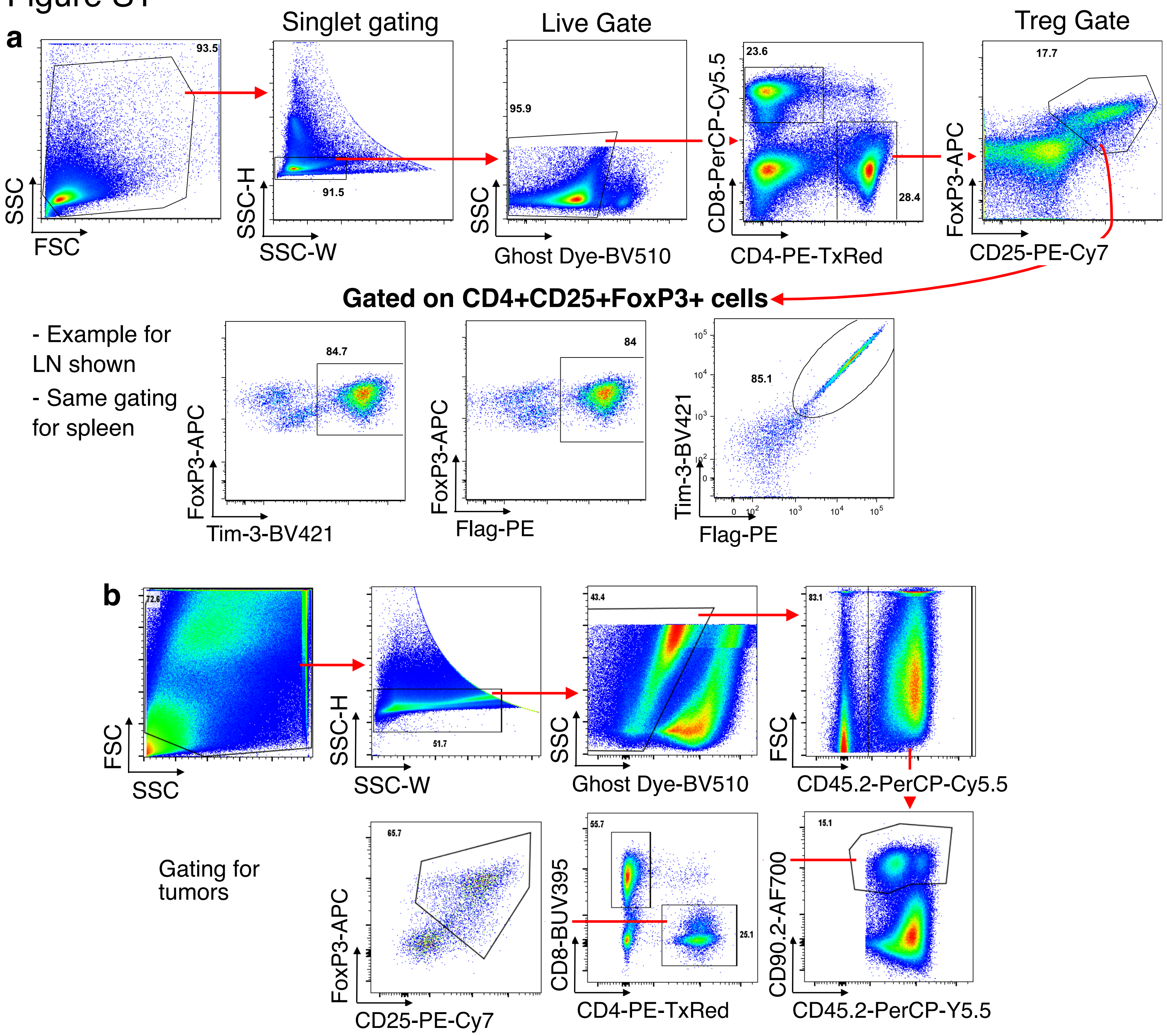


**Figure S1. Representative gating used for flow cytometry analysis of Treg from lymph node and spleen (panel a) or tumors (panel b).**

Figure S2

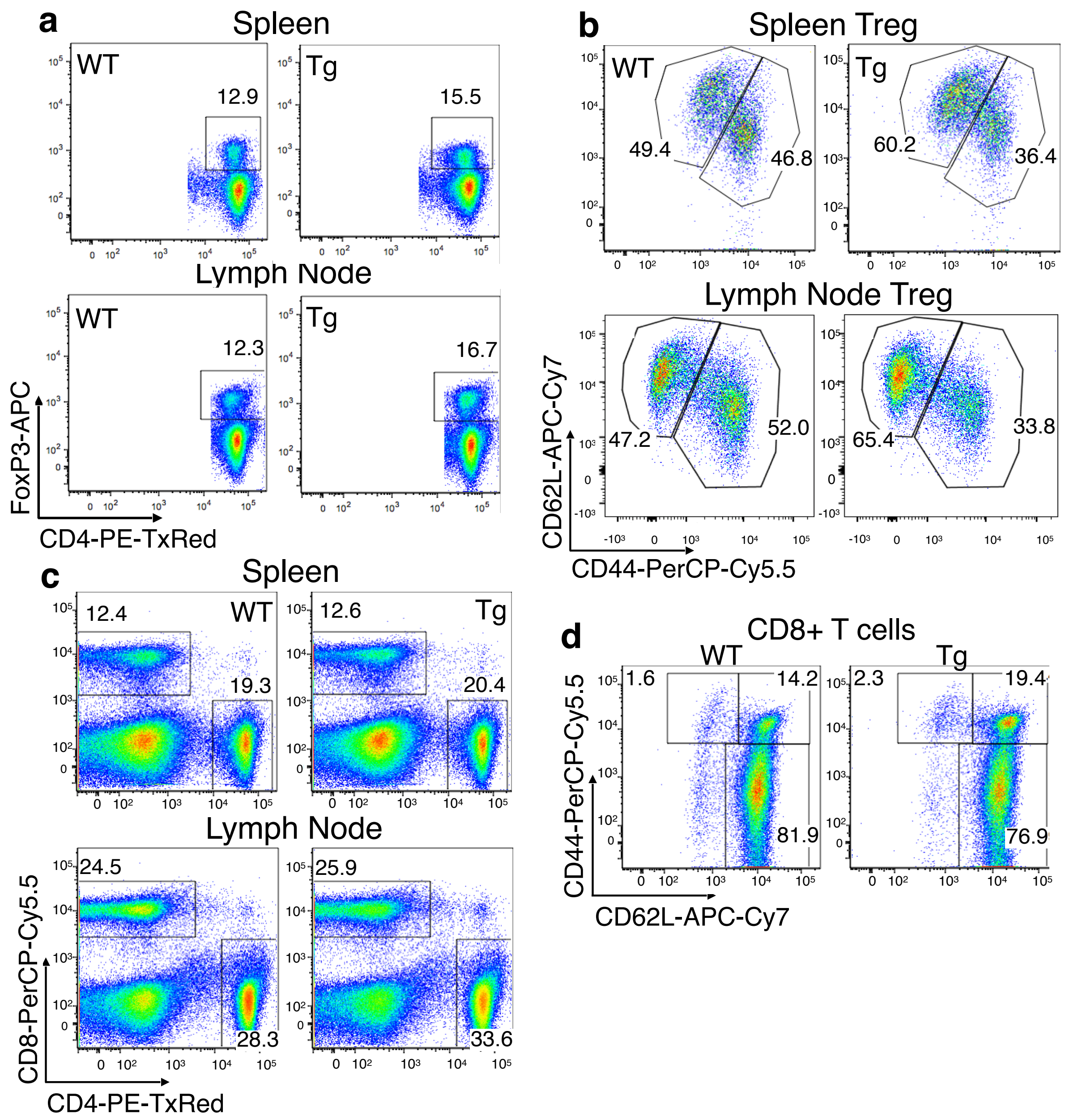


**Figure S2. Phenotype of peripheral conventional T cells and Treg from FoxP3-YFP-Cre x FSF-Tim-3 mice.** (a) Representative FoxP3 staining in spleen and lymph node of WT (Cre-only) and Tg animals. (b) Representative Treg activation phenotype in spleen and lymph node. (c) CD4/CD8 T cell compartments in spleen and lymph node of WT vs. Tg animals. (d) Activation status of conventional CD8+ T cells from spleen of WT vs. Tg animals.

### Figure S3

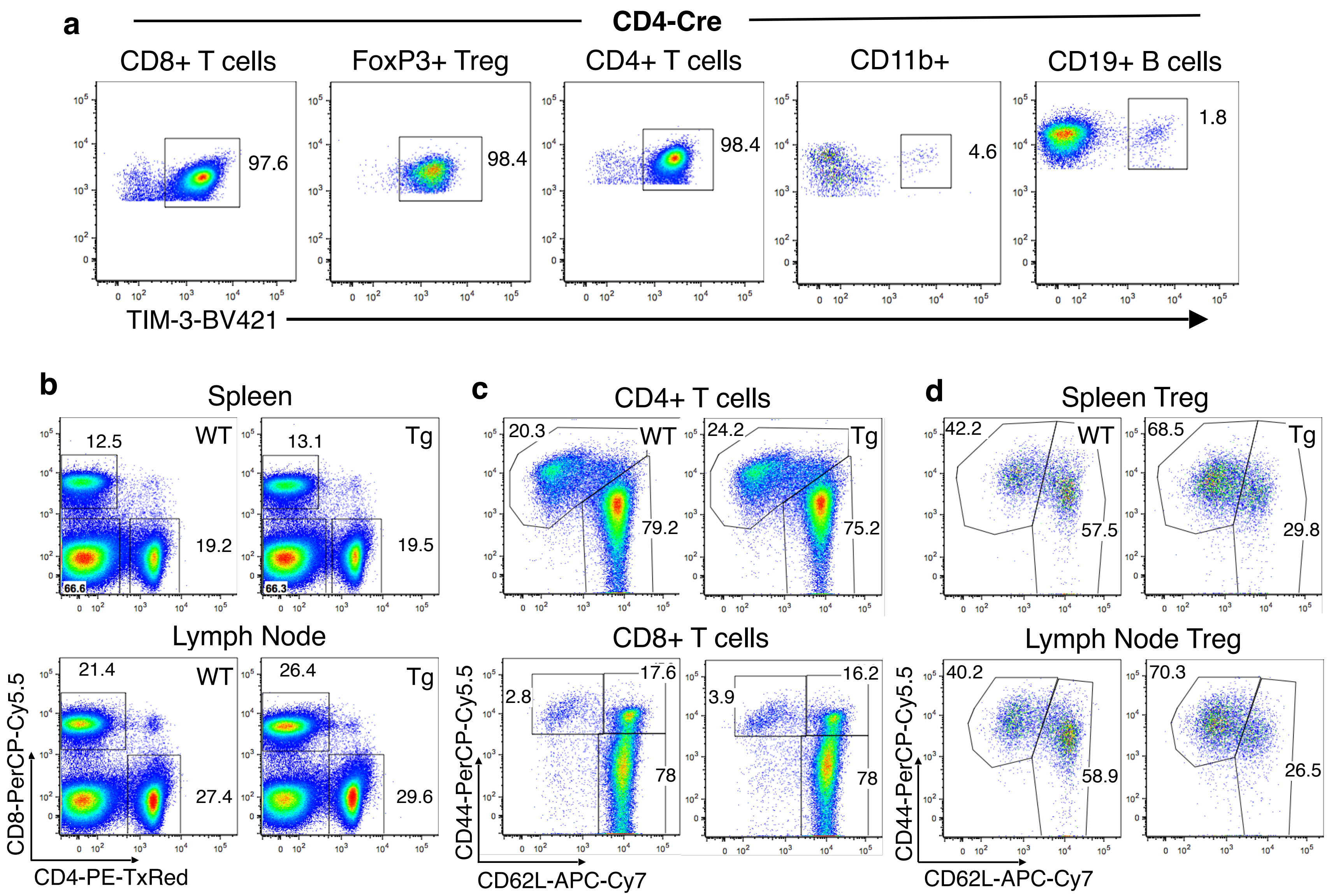


**Figure S3. Expression pattern and T cell phenotype in mice with Tim-3 expression driven by CD4-Cre.** (a) Faithful expression of the transgenic Tim-3 on mature T cells and Treg, but not myeloid or B cells. (b) Normal proportions of CD4+ and CD8+ T cells in spleen and lymph node. (c) Normal activation status of splenic conventional CD4+ and CD8+ T cells. (d) Increased activation of Treg in Tim-3 Tg animals with CD4-Cre.

Figure S4

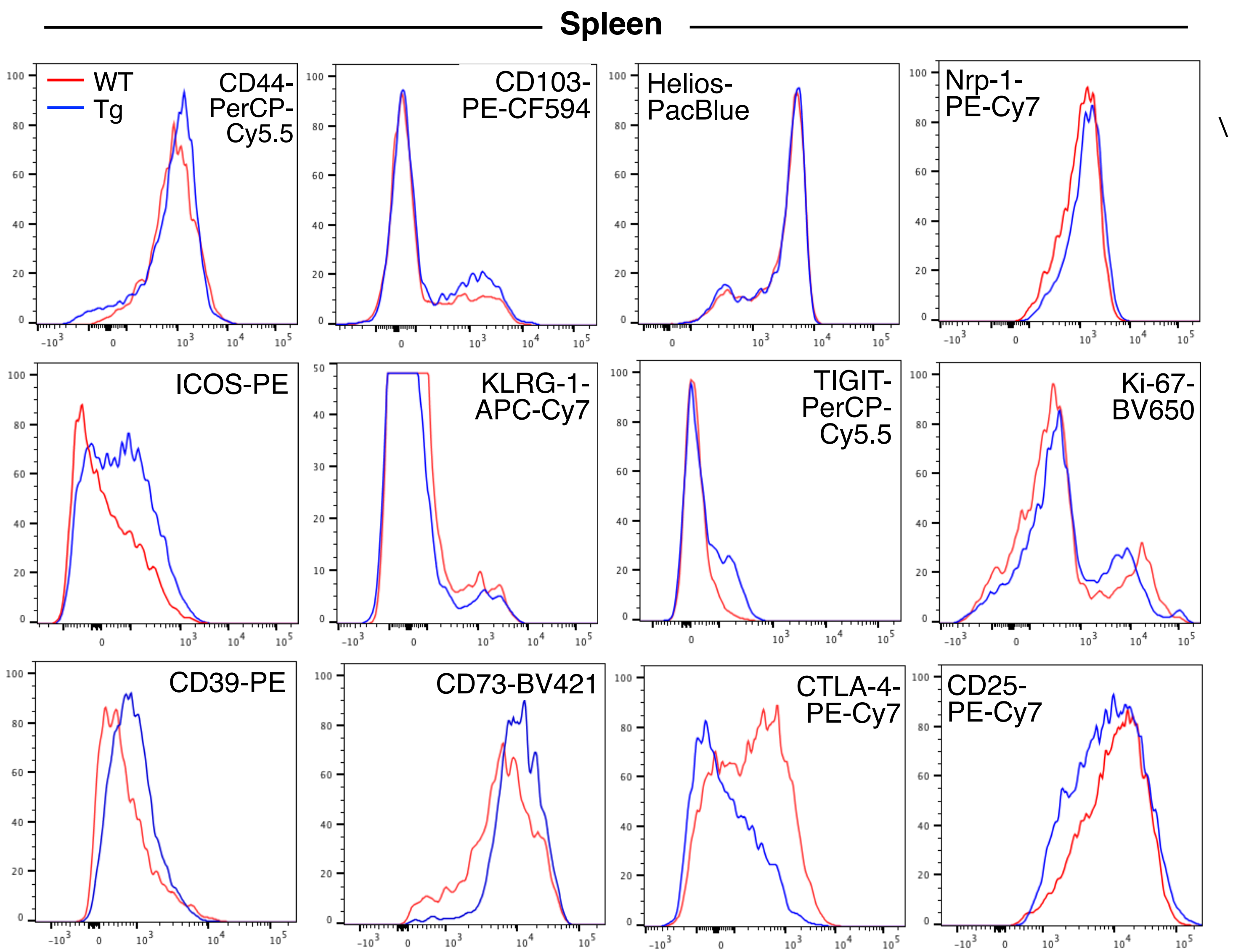


**Figure S4. Representative flow cytometry staining of multiple cell-surface markers on spleen Treg from FoxP3-eGFP-Cre-ERT2 x FSF-Tim-3 mice.** Mice expressing Cre only served at WT controls.

### Figure S5

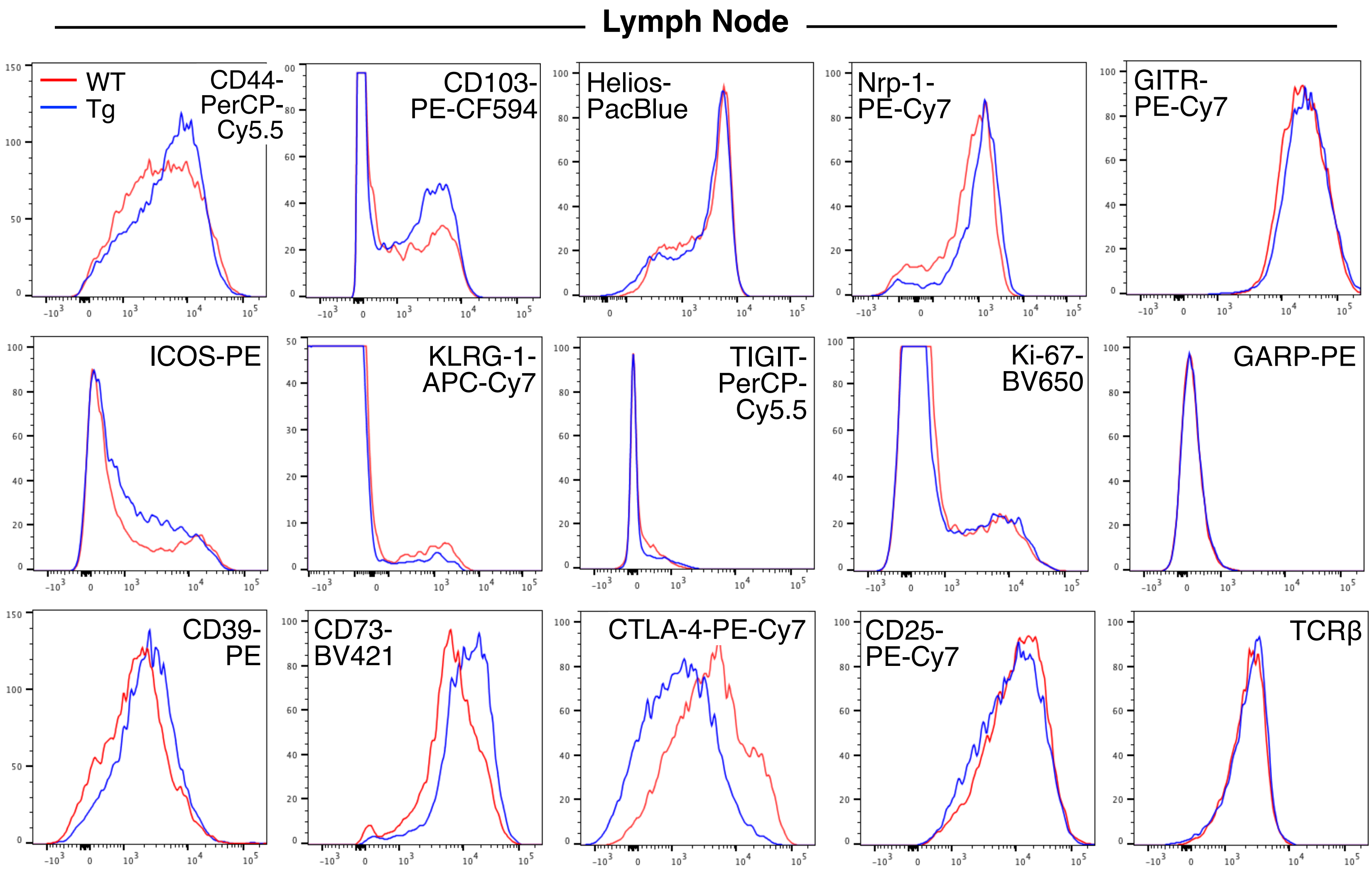


**Figure S5. Representative flow cytometry staining of multiple cell-surface markers on lymph node Treg from FoxP3-eGFP-Cre-ERT2 x FSF-Tim-3 mice.** Mice expressing Cre only served at WT controls.

### Figure S6

| Cytokines & Receptors | FSF-Tim3 $GFP^{Hi}$ | FSF-Tim3 $GFP^{Lo}$ |
|---|---|---|
| Il10 | 1884.5443 | 1746.1963 |
| Tgfb3 | 38.8542 | 709.176 |
| Il1rl2 | 20.1122 | 151.1671 |
| Il17rb | 12.8171 | 11.4716 |
| Il10ra | 7.2602 | 3.8906 |
| Il21 | 4.7899 | 28.443 |
| Il21r | 3.8637 | 3.7581 |
| Il12rb1 | 3.1383 | 1.1892 |
| Tgfb1 | 2.9897 | 1.7411 |
| Tnfrsf1a | 2.639 | 4 |
| Il27ra | 2.4794 | 2.8284 |
| Il2rb | 2.2346 | 0.5625 |
| Tnfrsf23 | 2.114 | 2.1435 |
| Il16 | 2.0562 | 2.2191 |
| Tnfrsf14 | 2.0139 | 2.5847 |
| Il2ra | 1.0281 | 0.0208 |
| Il20rb | 0.9659 | 1.0425 |
| Tnfsf13b | 0.895 | 0.4383 |
| Tnfsf8 | 0.8526 | 0.5783 |
| Tnfrsf12a | 0.8066 | 1.1892 |
| Tgfbr3 | 0.7955 | 3.5308 |
| Tnfsf10 | 0.7955 | 0.3231 |
| Tnfrsf8 | 0.722 | 0.0675 |
| Il1r2 | 0.7022 | 0.0621 |
| Tgfbr2 | 0.6643 | 1.3851 |
| Il1rap | 0.551 | 0.551 |
| Tnfrsf4 | 0.4665 | 0.0638 |
| Il18 | 0.4633 | 0.3253 |
| Tnfsf4 | 0.4633 | 0.1869 |
| Tnfrsf18 | 0.4601 | 0.1387 |
| Il18bp | 0.4234 | 0.2415 |
| Tgfbr1 | 0.4147 | 0.1582 |
| Tnfsf11 | 0.395 | 0.1426 |
| Il2 | 0.2624 | 1.2226 |
| Il4 | 0.183 | 0.2912 |

| Transcription Factors | FSF-Tim3 $GFP^{Hi}$ | FSF-Tim3 $GFP^{Lo}$ |
|---|---|---|
| Eomes | 14.1232 | 49.18 |
| Myc | 3.5554 | 1.9319 |
| Relb | 3.3636 | 2.1435 |
| Tbx21 | 3.0525 | 8.5742 |
| Gata1 | 2.9079 | 0.8706 |
| RelA | 2.4623 | 2.6027 |
| Tcf3 | 2.4453 | 2.7321 |
| Nr4a1 | 2.395 | 1.0943 |
| Klf6 | 2.2191 | 2.2191 |
| Ikzf3 | 2 | 1.6818 |
| Klf16 | 1.9862 | 1.6133 |
| Runx2 | 1.9725 | 1.1975 |
| Foxp3 | 1.9319 | 0.0854 |
| Runx3 | 1.7053 | 4.1989 |
| Lef1 | 1.5801 | 4.5631 |
| Ikzf4 | 1.5052 | 0.1571 |
| Prdm1 | 1.4743 | 1.2397 |
| Nfkbiz | 0.7684 | 0.409 |
| Klf11 | 0.727 | 0.6878 |
| Foxo4 | 0.722 | 1 |
| Hif1a | 0.712 | 0.5783 |
| Batf | 0.6643 | 0.4147 |
| Foxo3 | 0.6417 | 0.6878 |
| Ikzf1 | 0.6113 | 1.2311 |
| Id2 | 0.5743 | 1.7171 |
| Bcl6 | 0.5359 | 0.683 |
| Klf2 | 0.4633 | 0.7474 |
| Id3 | 0.4263 | 0.2892 |
| Nr4a3 | 0.4263 | 0.125 |
| Myb | 0.395 | 0.4175 |
| Nr4a2 | 0.3686 | 0.2698 |
| Ikzf5 | 0.2774 | 0.3015 |
| Ikzf2 | 0.2606 | 0.0467 |

| Ribosomal Factors | FSF-Tim3 $GFP^{Hi}$ | FSF-Tim3 $GFP^{Lo}$ |
|---|---|---|
| Rpn2 | 2.6759 | 2.7132 |
| Rprd1b | 1.8661 | 2 |
| Rpn1 | 1.7053 | 1.5157 |
| Rpl3 | 1.1329 | 1.0867 |
| Rprd2 | 1.021 | 0.9931 |
| Rpp14 | 0.6373 | 0.6462 |
| Rpp25l | 0.6113 | 0.6071 |
| Rpl7l1 | 0.6071 | 0.5141 |
| Rprd1a | 0.5946 | 0.6029 |
| Rpsa | 0.5434 | 0.5141 |
| Rpl4 | 0.5322 | 0.5249 |
| Rplp0 | 0.5212 | 0.5359 |
| Rpp40 | 0.4633 | 0.5285 |
| Rps4l | 0.4475 | 0.2398 |
| Rps2 | 0.4323 | 0.3057 |
| Rps3 | 0.4293 | 0.4293 |
| Rpl8 | 0.4175 | 0.4601 |
| Rpl13a | 0.409 | 0.3978 |
| Rpl7a | 0.4061 | 0.3121 |
| Rpl18a | 0.3585 | 0.3463 |
| Rpl29 | 0.3487 | 0.3099 |
| Rpp30 | 0.3345 | 0.3078 |
| Rpl28 | 0.3209 | 0.2774 |
| Rpl6 | 0.3164 | 0.2813 |
| Rps10 | 0.3143 | 0.2872 |
| Rpl10a | 0.3078 | 0.2755 |
| Rpl5 | 0.3057 | 0.2952 |
| Rpl14 | 0.2994 | 0.3392 |
| Rpl19 | 0.2994 | 0.2517 |

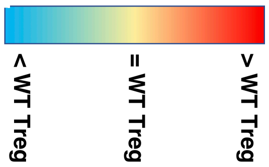


| Ribosomal, cont'd | FSF-Tim3 $GFP^{Hi}$ | FSF-Tim3 $GFP^{Lo}$ |
|---|---|---|
| Rps7 | 0.2045 | 0.2017 |
| Rps28 | 0.2017 | 0.2031 |
| Rpl34 | 0.1975 | 0.1756 |
| Rps19 | 0.1961 | 0.2003 |
| Rpl32 | 0.1948 | 0.1743 |
| Rplp1 | 0.1948 | 0.2045 |
| Rpl36 | 0.1934 | 0.1684 |
| Rps24 | 0.1921 | 0.1989 |
| Rpl26 | 0.1908 | 0.1684 |
| Rps27a | 0.1908 | 0.1882 |
| Rpl17 | 0.1895 | 0.1081 |
| Rpl31 | 0.1895 | 0.2003 |
| Rpl15 | 0.1882 | 0.1975 |
| Rps17 | 0.1882 | 0.1805 |
| Rpl41 | 0.1869 | 0.1684 |
| Rps21 | 0.1817 | 0.1882 |
| Rpl22 | 0.178 | 0.1539 |
| Rpl27 | 0.178 | 0.1593 |
| Rpl37rt | 0.1731 | 0.1426 |
| Rpl30 | 0.1719 | 0.1456 |
| Rpl23a | 0.1696 | 0.1416 |
| Rpl35 | 0.1696 | 0.1731 |
| Rpl37 | 0.1696 | 0.1406 |
| Rps14 | 0.1672 | 0.1604 |
| Rpl38 | 0.1661 | 0.134 |
| Rps23 | 0.1661 | 0.1661 |
| Rpl36a | 0.1649 | 0.1528 |
| Rpl39 | 0.1638 | 0.1368 |
| Rps27l | 0.156 | 0.1134 |
| Rpl32l | 0.155 | 0.1267 |
| Rpl35a | 0.1539 | 0.1571 |
| Rps27 | 0.1518 | 0.178 |
| Rps15a | 0.1456 | 0.1216 |
| Rps27rt | 0.1456 | 0.1708 |
| Rpl22l1 | 0.133 | 0.1358 |

**Figure S6. Relative expression of selected genes in Tim-3 Tg Treg ($FoxP3^{Hi}$ or $FoxP3^{Lo}$), compared with WT Treg.** Values are calculated based on average expression of Tim-3 Tg Treg populations, divided by average expression in FoxP3-Cre-only Treg, as determined by bulk RNAseq of the indicated sorted populations.

### Figure S7

| OxPhos | GFPHi Fold | GFPLo Fold |
|---|---|---|
| Atp5a1 | 1.8 | 1.7 |
| Atp5b | 2 | 1.9 |
| Atp5c1 | 0.37 | 1.9 |
| Atp5d | 1 | 0.69 |
| Atp5e | 0.18 | 0.16 |
| Atp5f1 | 0.34 | 0.36 |
| Atp5g1 | 0.24 | 0.18 |
| Atp5g2 | 0.31 | 0.26 |
| Atp5g3 | 0.33 | 0.31 |
| Atp5h | 0.24 | 0.26 |
| Atp5j | 0.18 | 0.16 |
| Atp5j2 | 0.17 | 0.15 |
| Atp5k | 0.17 | 0.15 |
| Atp5l | 0.16 | 0.16 |
| Atp5o | 0.26 | 0.23 |
| Atp5s | 0.67 | 0.51 |
| Atp6ap1 | 1.8 | 2.5 |
| Atp6ap2 | 0.86 | 1.3 |
| Atp6v0a1 | 4.3 | 1 |
| Atp6v0a2 | 2.8 | 4.8 |
| Atp6V0b | 0.58 | 0.63 |
| Atp6V0c | 1.4 | 1.7 |
| Atp6V1a | 0.68 | 1 |
| Atp6V1b2 | 2.2 | 2.4 |
| Atp6V1c1 | 1 | 1 |
| Atp6V1d | 0.41 | 0.5 |
| Atp6V1e1 | 0.45 | 0.49 |
| Atp6V1f | 0.25 | 0.25 |
| Atp6V1g1 | 0.34 | 0.43 |
| Atp6V1g2 | 1 | 1 |
| Atp6V1g3 | 0.14 | 0.04 |
| Atp6V1h | 1 | 1.4 |
| Atp6Vod1 | 1 | 1 |
| Atp6Voe | 0.24 | 0.29 |
| Cox10 | 2 | 2.8 |
| Cox11 | 4.2 | 4.4 |
| Cox14 | 0.24 | 0.26 |
| Cox15 | 3.7 | 3.5 |
| Cox16 | 0.24 | 0.29 |
| Cox17 | 0.27 | 0.24 |
| Cox18 | 1 | 1 |
| Cox19 | 0.52 | 0.56 |
| Cox20b | 0.21 | 0.17 |
| Cox4i1 | 0.28 | 0.26 |
| Cox5a | 0.29 | 0.27 |
| Cox5b | 0.23 | 0.21 |
| Cox6a1 | 0.23 | 0.27 |
| Cox6b1 | 0.23 | 0.2 |
| Cox6c | 0.15 | 0.13 |
| Cox7a2 | 0.16 | 0.14 |
| Cox7b | 0.17 | 0.14 |
| Cox7c | 0.17 | 0.15 |
| Cox8a | 0.26 | 0.27 |
| Cyc1 | 1.3 | 1.2 |

| OxPhos | GFPHi Fold | GFPLo Fold |
|---|---|---|
| Lhpp | 2.70 | 2.8 |
| mt-Atp6 | 0.19 | 0.18 |
| mt-Co1 | 0.4005 | 0.4061 |
| mt-Co2 | 0.1539 | 0.1518 |
| mt-Co3 | 0.1051 | 0.1066 |
| mt-Nd1 | 0.22 | 0.24 |
| mt-Nd2 | 0.26 | 0.27 |
| mt-Nd4 | 0.34 | 0.33 |
| mt-Nd5 | 0.56 | 0.51 |
| mt-Nd6 | 0.3 | 0.27 |
| Ndufa10 | 0.64 | 0.74 |
| Ndufa11 | 0.23 | 0.22 |
| Ndufa2 | 0.17 | 0.18 |
| Ndufa3 | 0.19 | 0.19 |
| Ndufa4 | 0.16 | 0.15 |
| Ndufa5 | 0.16 | 0.13 |
| Ndufa6 | 0.19 | 0.18 |
| Ndufa7 | 0.21 | 0.2 |
| Ndufa8 | 0.33 | 0.34 |
| Ndufb10 | 0.26 | 0.29 |
| Ndufb2 | 0.22 | 0.24 |
| Ndufb5 | 0.21 | 0.2 |
| Ndufb6 | 0.21 | 0.17 |
| Ndufb7 | 0.25 | 0.21 |
| Ndufb8 | 0.22 | 0.19 |
| Ndufb9 | 0.26 | 0.21 |
| Ndufc1 | 0.18 | 0.18 |
| Ndufc2 | 0.22 | 0.18 |
| Ndufs1 | 1.27 | 1.4 |
| Ndufs2 | 1.4 | 1.4 |
| Ndufs3 | 0.53 | 0.54 |
| Ndufs4 | 0.21 | 0.22 |
| Ndufs5 | 0.21 | 0.2 |
| Ndufs6 | 0.21 | 0.19 |
| Ndufs7 | 0.4 | 0.41 |
| Ndufs8 | 0.36 | 0.28 |
| Ndufv1 | [illegible] | 1.3 |
| Ndufv2 | [illegible] | 0.29 |
| Ndufv3 | [illegible] | 0.24 |

> WT Treg
= WT Treg
< WT Treg

| Glycolysis | GFPHi Fold | GFPLo Fold |
|---|---|---|
| Acss1 | 1 | 1 |
| Acss2 | 2.9 | 6.4 |
| Acyp1 | 0.21 | 0.23 |
| Adh1 | 3 | 20.1 |
| Adh5 | 0.67 | 0.78 |
| Akr1a1 | 1 | 1 |
| Aldh1b1 | 8.1 | 6.9 |
| Aldh2 | 2.8 | 4.5 |
| Aldh3a2 | 3 | 3.5 |
| Aldh7a1 | 3.2 | 1 |
| Aldh9a1 | 1.9 | 2.2 |
| Aldoa | 2.6 | 2.9 |
| Aldoc | 3.4 | 1 |
| Bpgm | 1.5 | 2.2 |
| Dlat | 1 | 1 |
| Dld | 1 | 1 |
| Eno1 | 4.8 | 4.8 |
| Eno2 | 1 | 1 |
| Eno3 | 1.5 | 0.5 |
| G6pc3 | 2.1 | 1 |
| Galm | 2 | 1 |
| Gapdh | 8.7 | 11.6 |
| Gpi1 | 2.9 | 2.5 |
| Hk1 | 3.5 | 3 |
| Hk2 | 1 | 0.08 |
| Ldha | 1 | 1 |
| Ldhb | 0.53 | 0.28 |
| Ldhd | 1 | 2.6 |
| Mdh1 | 0.38 | 0.34 |
| Mdh2 | 1 | 1 |
| Mpc1 | 0.3 | 0.27 |
| Mpc2 | 0.24 | 0.24 |
| Pck2 | 3.8 | 2.4 |
| Pdha1 | 1.4 | 1.4 |
| Pdhb | 0.72 | 0.8 |
| Pdhx | 1 | 1 |
| Pdk1 | 1 | 3 |
| Pdk2 | 5.1 | 3.4 |
| Pdk3 | 1 | 0.6 |
| Pdp1 | 1 | 1 |
| Pdp2 | 1.8 | 1.8 |
| Pdpr | 5.2 | 4.8 |
| Pfkl | 9.1 | 10.3 |
| Pfkm | 4 | 3.2 |
| Pfkp | 4.5 | 5.7 |
| Pgam1 | 1 | 1 |
| Pgam5 | 3.5 | 3.5 |
| Pgk1 | 1 | 1 |
| Pgm1 | 3.7 | 3.7 |
| Pgm3 | 1 | 1 |
| Pkm | 7.1 | 7.1 |
| Slc2a1 | 2.2 | 2.6 |
| Slc2a3 | 1 | 1 |
| Tpi1 | 1.9 | 1 |

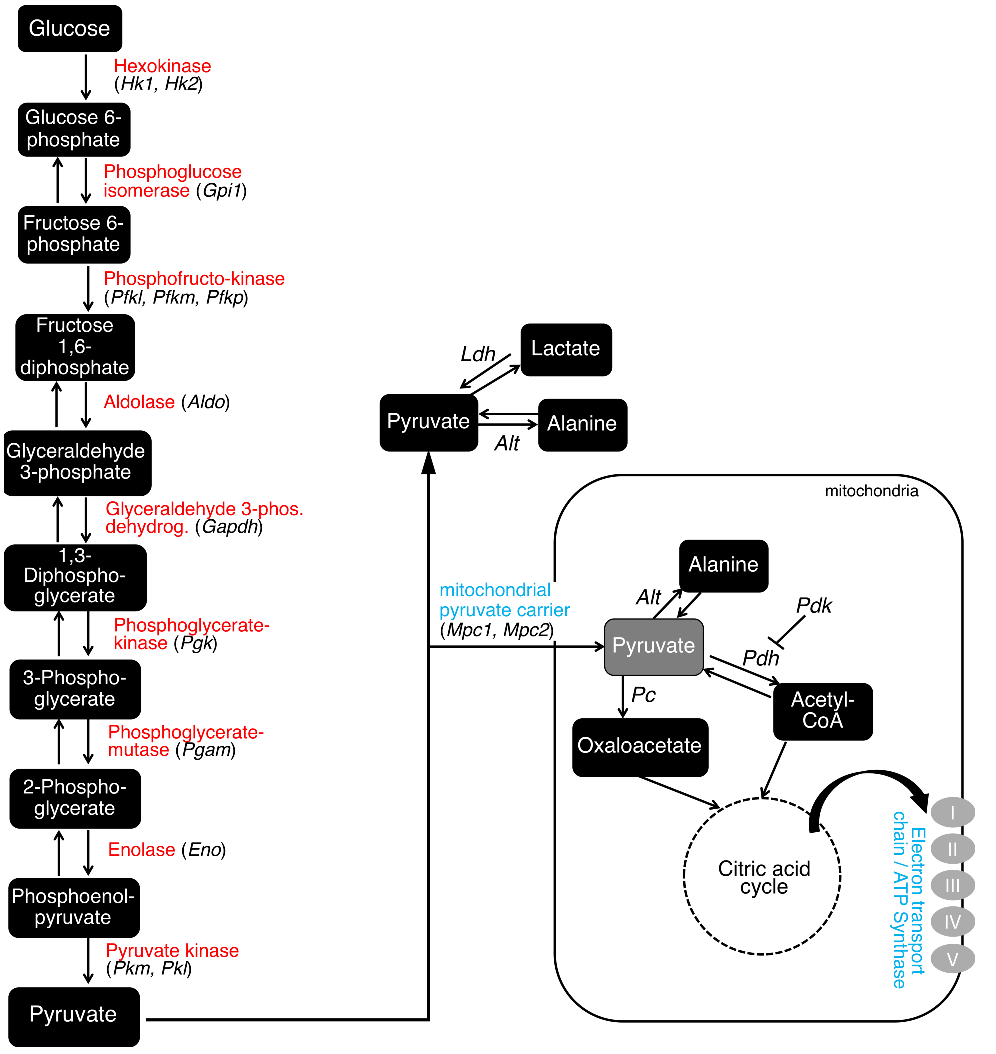


**Figure S7. Broad effects of Tim-3 expression on Treg metabolic gene signatures.** Heat maps represent relative expression of indicated glycolysis and oxidative phosphorylation/ mitochondrial electron transport chain gene, in FoxP3-eGFP$^{Hi}$ and FoxP3-eGFP$^{Lo}$ Treg, compared with FoxP3-eGFP-Cre only control Treg. Right – schematic overview of glycolysis and ox phos pathways highlighting nodes upregulated (red) or downregulated (blue) in Tim-3 Tg Treg.

### Figure S8

| | | | | |
|---|---|---|---|---|
| HAVCR2 | PRF1 | PHLDA1 | CCR1 | RBPJ |
| TNFRSF18 | ENO1 | CCL4 | IFNG | TMBIM6 |
| LAG3 | IL1R2 | CD82 | TUBB | VCP |
| GAPDH | GZMH | HLA-A | CSF1 | CALR |
| CTSC | ICOS | IL2RG | PDCD1 | NDUFC2 |
| GZMB | TPI1 | CXCL13 | DNPH1 | CXCR3 |
| CCL5 | PTMS | RANBP1 | FKBP1A | CKLF |
| GNLY | CCND2 | KLRD1 | PMVK | UCP2 |
| CST7 | GZMA | SPOCK2 | ISG15 | LDHA |
| BST2 | MYL6 | CTSW | LAPTM4B | EFHD2 |
| CD7 | TNFRSF1B | COX5A | SOX4 | ZBTB32 |
| TNFRSF4 | LY6E | IFITM2 | MT2A | ACTB |
| LAIR2 | PGAM1 | CD8A | SLC27A2 | CCL4L2 |
| NKG7 | CXCR6 | CD2 | MAP2K3 | ATP5MC3 |
| CTLA4 | APOBEC3G | GBP2 | CD3D | TIGIT |
| PKM | SH2D2A | CCL3 | PGK 1 | MX1 |
| IFI6 | LGALS1 | LINC0187 1 | IL2RA | OASL |
| ACP5 | BATF | ATOX1 | S100A4 | GADD45A |
| TYMP | SYNGR2 | TNFRSF9 | TXN | SERPINB9 |
| ID2 | CD63 | IL2RB | DUSP4 | COX7A2 |

**Figure S8. List of differentially expressed genes between TIM-3+ and TIM-3- Treg in human HNC TIL scRNAseq dataset used for KEGG pathway analysis.**
